# Single cell multi-omics enables high-resolution identification and functional purification of human acute myeloid leukemia stem cells

**DOI:** 10.64898/2026.07.12.737989

**Authors:** Asiri Ediriwickrema, Yusuke Nakauchi, Thomas Köhnke, Amy C. Fan, Xiaoyi Hu, Brooks A. Benard, Daiki Karigane, Miles H. Linde, Aaron M. Newman, Andrew J. Gentles, Ravindra Majeti

**Author notes:** Corresponding author; address: Lokey Stem Cell Research Building, Room G3167A, 265 Campus Dr, Stanford, CA 94305, USA.

## Abstract

In human acute myeloid leukemia (AML), a sub-population of leukemia stem cells (LSCs) drive disease initiation, therapeutic resistance, and relapse. However, the lack of reliable markers to distinguish LSCs from bulk leukemia cells has impeded progress in studying LSC pathogenesis and developing meaningful LSC-specific diagnostics and therapeutics. Existing LSC gene signatures, derived from bulk populations, cannot definitively identify LSCs at single-cell resolution. To address this, we analyzed large patient cohorts with bulk gene expression data and single-cell multi-omic assays to identify a prognostic gene signature that is specifically enriched in a clinically adverse AML sub-population. Using this signature, we defined and prospectively isolated CD34+CD90-CLL1-CD69+CD53- immunophenotypic LSCs that are significantly enriched for LSC content based on limiting dilution xenotransplantation assays. Our findings demonstrate the power of single-cell multi-omics to precisely identify a clinically relevant LSC population and establish a clear framework for future translational research in AML.

**Key Points:**

- Single cell multi-omics identifies human AML LSCs at high resolution.
- *HOPX* and *SOCS2* co-expression (hrLSC2) defines a prognostic gene signature in de novo acute myeloid leukemia.
- hrLSC2 marks an AML subpopulation (iLSCs) with a distinct immunophenotype.
- iLSCs can be purified using flow cytometry and are significantly enriched for LSCs.

## Introduction

Acute myeloid leukemia (AML) is an aggressive bone marrow cancer that is associated with a five year survival rate of less than 32%^1^. Most patients are either refractory to treatment or relapse with incurable disease^2^. Although current risk stratification systems can provide guidance on treatment^3,4^, they rely primarily on genetic factors which may incompletely characterize AML severity^5^. The cancer stem cell model can provide additional prognostic insights by describing the functional heterogeneity of the disease cells^6^. According to the cancer stem cell model, AML is organized as a cellular hierarchy that develops from the leukemic transformation of hematopoietic stem and progenitor cells (HSPCs) into leukemia stem cells (LSCs)^5,7^. LSCs are defined as cells capable of transplanting leukemia into immunodeficient mice and higher LSC frequency is associated with treatment resistance and disease relapse^6,7^. Similarly, high LSC content determined by gene expression signatures derived from LSC-enriched cellular fractions of patient samples are associated with poor clinical outcomes^8–10^. These observations suggest that LSCs may need to be eliminated to cure patients. A significant limitation of these prior studies is that the corresponding LSC gene signatures were derived from populations of leukemic cells enriched for LSCs, where the frequency of functional LSCs is variable and heterogeneous. Importantly, it is unclear if these signatures define a unique LSC population or a coordinated gene expression program across a mixture of cells. Therefore, AML LSCs remain ill-defined at the single cell level, which has impaired our ability to study their biological and clinical relevance and advance LSC-targeted therapies.

Effectively addressing these limitations requires the prospective isolation of LSCs at high purity using surface markers. Although there has been a long history of defining LSC-enriched sub-populations in this way^11–16^, a highly specific immunophenotype remains elusive as it varies significantly from patient to patient. Initially, AML LSC sub-populations were identified in the CD34+CD38- compartment through xenotransplantation assays^17^, and were reported to lack the canonical hematopoietic stem cell (HSC) marker CD90^12^. Subsequent studies have demonstrated that LSCs are present in both CD34+CD38+ and CD34- fractions at lower frequencies^9–11,18,19^. Over the following years, several groups have identified additional candidate LSC antigens including TIM3^15,20^, CD99^16^, CD123^21,22^, CD44^23^, CLL1^24^, CD25^25^, CD96^26^, CD47^14^, GPR56^27^, CD93^28^, CD244^29^, S1PR3^30^, and CD117^29^. Despite these extensive analyses, highly specific AML LSC markers have not been identified. Part of the difficulty is due to different standards for defining LSCs as only a few studies performed limiting dilution assays (LDAs) to quantify LSC content, and different xenograft models can influence transplantation efficiency and stem cell quantification^31–33^. Amongst the xenotransplantation studies that performed LDAs, Eppert et al. observed an LSC frequency between 1/1,600–1/1,100,000 cells in CD34+CD38- cells from 16 primary samples^9^, Xie et al. observed an LSC frequency of 1/882–1/33,314 in CD34+CD33+S1PR3low/- cells from 3 primary samples^30^, and Pabst et al. observed an LSC frequency of 1/519 in CD34+GPR56+ albeit in a single AML case^27^. The lack of a specific and generalizable LSC immunophenotype and inability to purify cells at the single cell level has impaired our ability to directly study AML LSC pathogenesis and connect LSC biology broadly with clinical outcomes. Specifically, it is not known if prognostic LSC gene signatures are highly expressed in single cells, if these cells express a unique immunophenotype, and if that immunophenotype can enrich for functional LSCs.

Here, we systematically address these limitations by basing our analysis on four fundamental principles (Figure 1A): 1) LSC burden is associated with worse clinical outcomes, 2) LSC signatures should be expressed in a cell type specific manner, 3) these cells can be purified by flow cytometry, and 4) LSCs should demonstrate a higher capacity to engraft leukemia in immunodeficient mice compared to bulk AML cell populations. Using these principles, we derived a two gene high risk LSC gene expression signature (i.e. hrLSC2) that is both strongly prognostic and highly expressed in specific AML cells. Cells with increased expression of hrLSC2 were defined as candidate LSCs (cLSCs) and were enriched in a population of CD34+CD90-CLL1-CD69+CD53- cells, i.e. immunophenotypic LSCs (iLSCs), based on concurrent scRNA-seq and surface marker quantification. iLSCs were prospectively isolated by Fluorescence-Activated Cell Sorting (FACS) and contained significantly greater LSC frequency compared to bulk CD34+ cells. Overall, our findings provide updated computational and flow cytometry-based strategies for isolating prognostically adverse LSCs in adult human AML.

**Figure 1:**
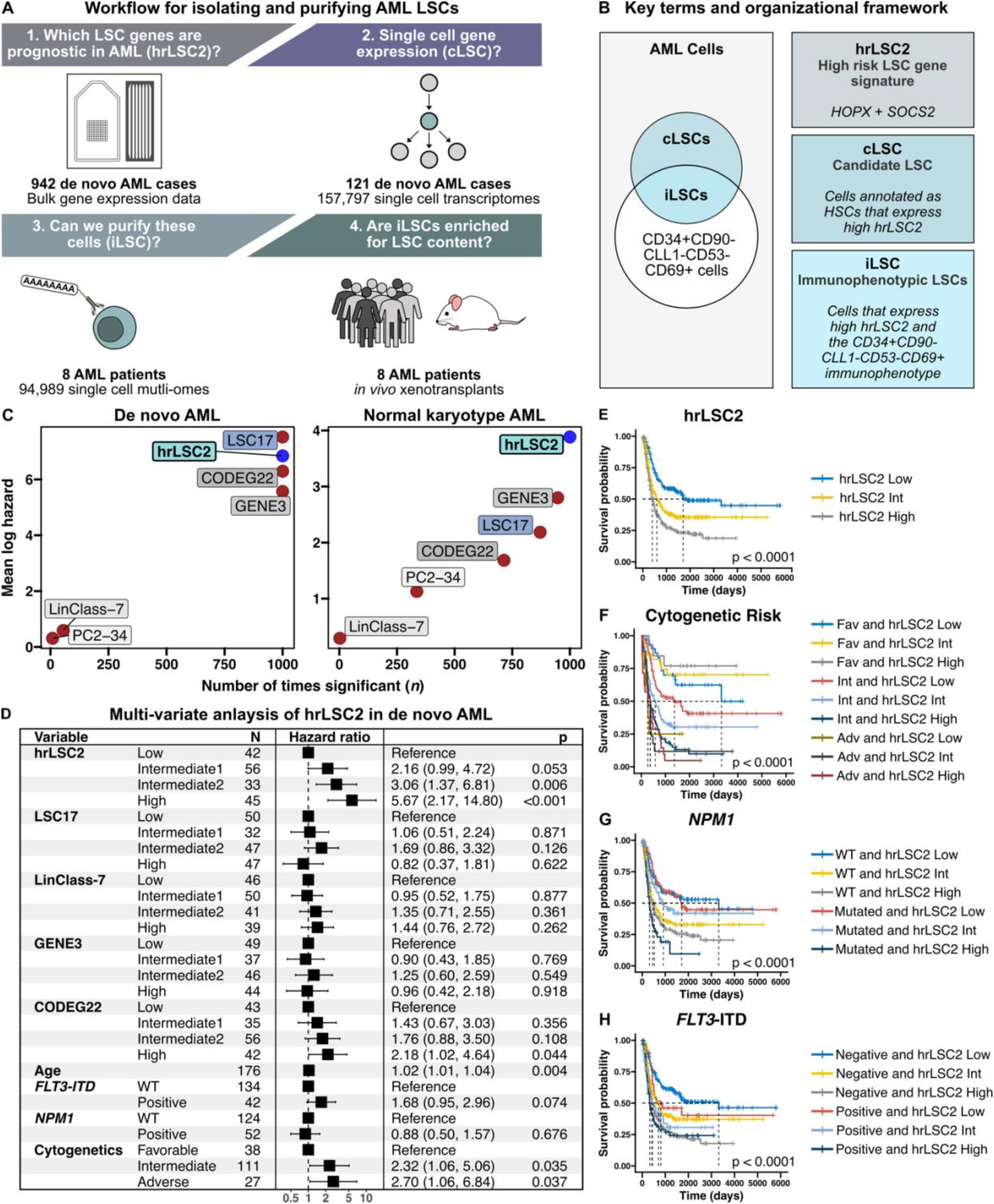
hrLSC2 is a prognostic gene signature in de novo AML. A) General workflow for identifying an LSC-specific gene signature in de novo AML. B) Key terminology used throughout the manuscript and Venn diagram illustrating relevant LSC sup-populations evaluated in this study. C) Results of iterative coxph regression analysis (*n*=1000) in de novo and normal karyotype AML evaluating the performance of hrLSC2 against previously published prognostic AML signatures for predicting survival. D) Multi-variate coxph survival analysis evaluating the association between prognostic AML signatures, age, high and low risk mutation status, and cytogenetic risk categories. E-H) Kaplan-Meier survival analysis evaluating hrLSC2 status in combination with established high and low risk features. Statistical significance was tested using either the Wald or Log-rank test.

### Definitions (Figure 1B)

1. hrLSC2 (high risk two gene LSC signature): LSC gene expression signature comprised of *HOPX* and *SOCS2*
2. cLSC (candidate LSC): AML cells annotated as HSCs using a normal HSPC-specific classifier that also highly express hrLSC2
3. iLSC (immunophenotypic LSC): CD34+CD90-CLL1-CD69+CD53- AML cells that express high hrLSC2 and are enriched for cLSCs and functional LSCs

## Results

### hrLSC2 is a simple yet highly prognostic LSC signature in de novo AML

AML risk stratification systems guide clinicians on the diagnosis, prognostication, and management of patients and rely primarily on molecular and cytogenetic profiling^3,4^. Using these systems, clinicians can categorize AML patients as either favorable, intermediate, or adverse risk which can predict prognosis and influence treatment. Most AML arises de novo, i.e. without prior exposure to chemoradiation or antecedent hematologic disorders and are categorized as intermediate risk. Within this majority, patients have highly variable responses to treatment and clinical outcomes. This highlights a significant limitation of current risk models which may benefit from including features of the AML hierarchy such as LSC content^5^. Bulk gene expression has been shown to correlate with LSC content; however, numerous studies have subsequently generated data that could refine the initial LSC signatures. We therefore revisited the task of identifying an LSC specific and prognostically relevant signature, by curating a large AML cohort (*n*=1,949) of published bulk gene expression data with survival annotations^34–40^. We focused our analysis on de novo AML evaluated at diagnosis and excluded acute promyelocytic leukemia which can be considered a separate AML subtype as it has a distinct treatment paradigm and highly favorable outcomes. This resulted in a core set of 942 cases (Table ST1, Figure S1). We then narrowed the gene expression search space to include genes that have been associated with LSC biology (Table ST2)^8–10,41^. After data normalization and scaling, the cohort was split into training (*n*=626) and test (*n*=316) sets which were well balanced (Figure S2). We subsequently evaluated the training dataset using ensemble machine learning methods to identify genes that were most associated with shorter survival. We aggregated the ranked lists (Table ST3) generated from these ensemble methods to identify the top features^42^, and observed that higher expression of *HOPX* and *SOCS2* were the most important features associated with worse survival in both de novo and normal karyotype AML, which was used as a surrogate for intermediate risk AML (Table ST4).

We then generated a weighted score based on *HOPX* and *SOCS2* expression, i.e. hrLSC2 (Figure 1B), using the training dataset and evaluated its performance for predicting survival in the test dataset by conducting iterative cox proportional-hazards (coxph) regressions (*n*=1,000; Figure 1C). hrLSC2 and LSC17^10^, a rigorously validated LSC signature, performed better than other gene signatures^43–45^ in de novo AML (Figure 1C, left panel). When evaluating normal karyotype AML, hrLSC2 was superior to other LSC signatures (Figure 1C, right panel). The improved performance of hrLSC2 was highlighted further in a multi-variate coxph analysis including other LSC signatures, age, sex, *FLT3*-ITD status (high risk mutation), *NPM1* mutation status (low risk mutation), and cytogenetic risk category (Figure 1D; *n*=176 test cases containing both mutation and cytogenetic annotations). Compared to all other covariates, hrLSC2 was most significantly associated with worse survival in de novo AML (Figure 1E) and further stratifies survival in conjunction with established risk features (Figure 1F-H). Overall, this analysis supports that hrLSC2 is highly prognostic and may improve patient risk stratification when incorporated with genomic features.

**Figure S1:**
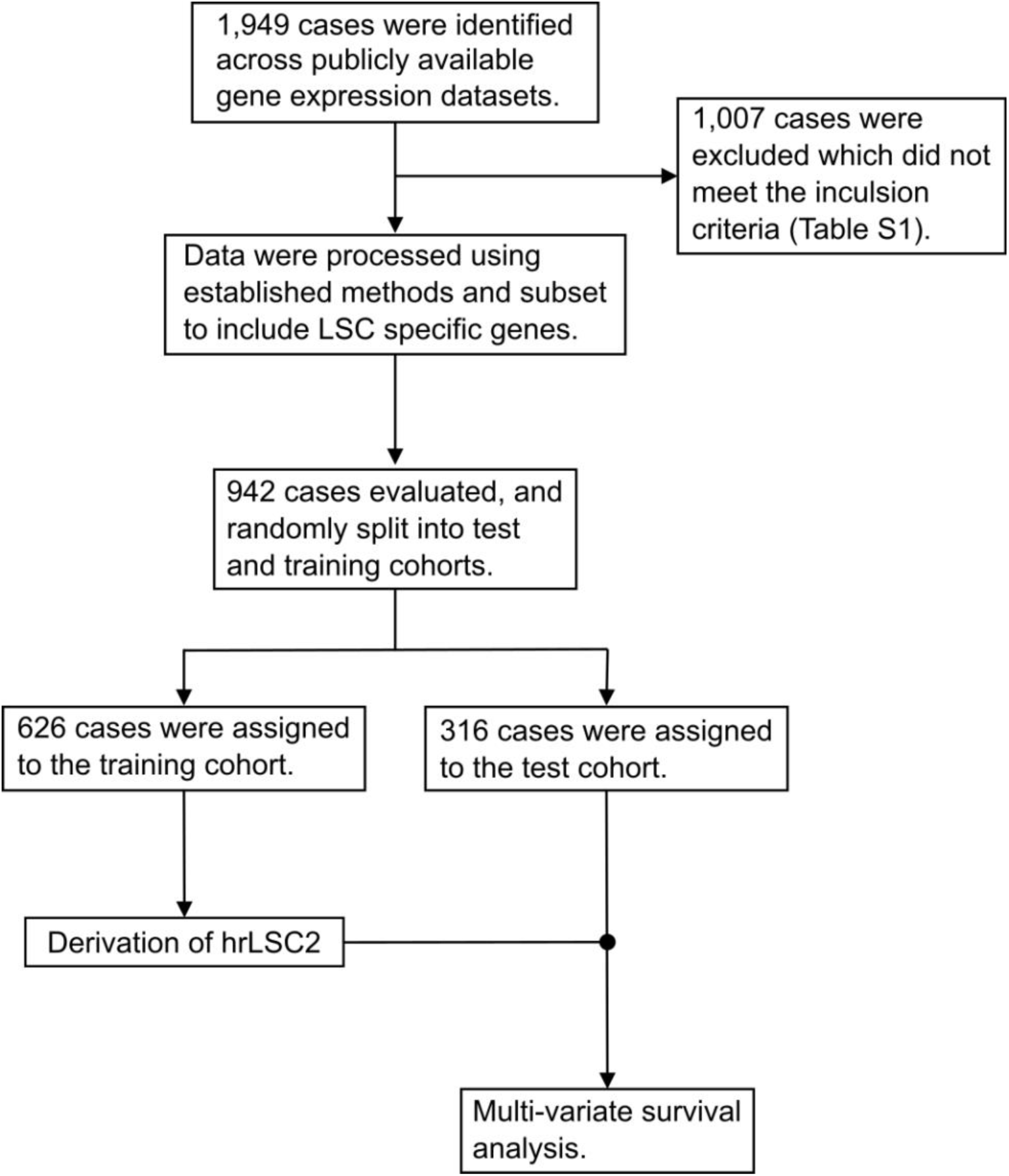
Dataset curation, filtration, and analysis workflow for deriving hrLSC2.

**Figure S2:**
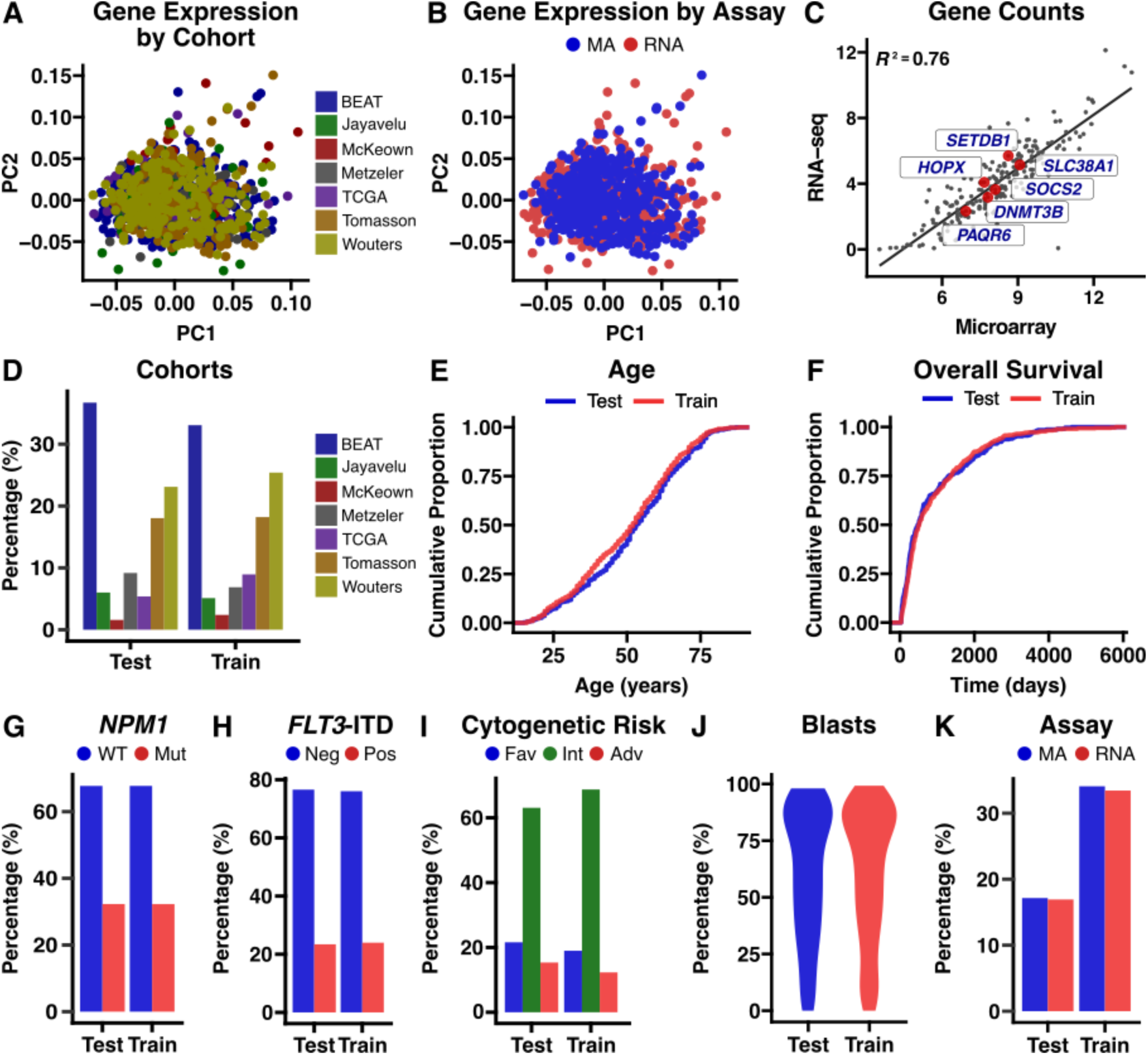
Description of key dataset metrics. Principal component plot of gene expression labeled by (A) cohort and (B) assay subtype. C) Correlation between processed gene counts from RNA-seq and MA data with highly prognostic genes labeled. D) Bar plot showing relative proportions of each cohort in the training and test datasets. Comparison of training and test dataset compositions for key characteristics including (E) age, (F) survival, (G-J) clinical features, and (K) assay. MA – microarray, Neg – negative, Pos – positive, RNA – RNA-seq.

### High hrLSC2 expression marks a prognostically relevant sub-population in de novo AML

We subsequently evaluated whether hrLSC2 expression could allow us to specifically identify a highly expressing cellular sub-population across a large cohort of de novo AML. To address this goal, we compiled publicly and internally generated scRNA-seq data (*n*=985,422 cells) from patients with de novo AML (*n*=121) and healthy donors (*n*=25) (Figure S3, Figure 2A, S4A, and Table ST7). After applying the same processing and quality control methods to each dataset (Figure S3), we removed lymphocytes and annotated cells using an HSPC-specific classifier that we previously developed to accurately annotate human HSPCs in scRNA-seq data^46^. The data (*n*=157,797 cells) were then integrated using scGen (Figure 2B)^47^ and hrLSC2 expression was calculated for each cell. To determine if hrLSC2 could identify an AML sub-population, we first evaluated its expression across all AML cells and observed a multi-modal distribution (Figure S4B). We then used the expectation-maximation algorithm to fit finite mixtures based on hrLSC2 expression^48^, which allowed us to identify cells that express hrLSC2 at high levels. These cells were enriched for cells mapping to the most primitive progenitors, including HSCs and LMPP1s (Figure S4C) and were more homogenous based on lower diversity indices (Figure S4D) and cellular spread (Figure S4E) compared to all cell types for each donor (Methods). The data were down sampled to ensure an equivalent number of cells were being compared to the hrLSC2 high sub-population. As expected, cells annotated as HSCs and LMPP1s not only expressed hrLSC2 at high levels (Figure 2C) but also contained a high fraction of cells that co-expressed *HOPX* and *SOCS2* compared to all other cell types (Figure 2C-D). These observations support that high hrLSC2 derived from both genes, rather than high levels from one gene only, is important for marking these cells. To computationally enrich for hrLSC2-expressing cells within the AML HSPC compartment, we applied an expectation-maximization algorithm to identify cells exhibiting high hrLSC2 expression. Since AML-HSCs demonstrated the highest hrLSC2 expression levels among all cellular subsets (Figure 2C) and is considered the most primitive cell type, we focused the subsequent analysis on this population. AML-HSCs expressing high hrLSC2 were designated as candidate LSCs (cLSCs) (Figure 2E) and showed significant enrichment for cells co-expressing *HOPX* and *SOCS2* relative to AML-HSCs with lower hrLSC2 expression. AML-HSCs exhibiting low hrLSC2 expression retained their original cellular annotation as an AML-HSC (Figure 2F-G).

**Figure 2:**
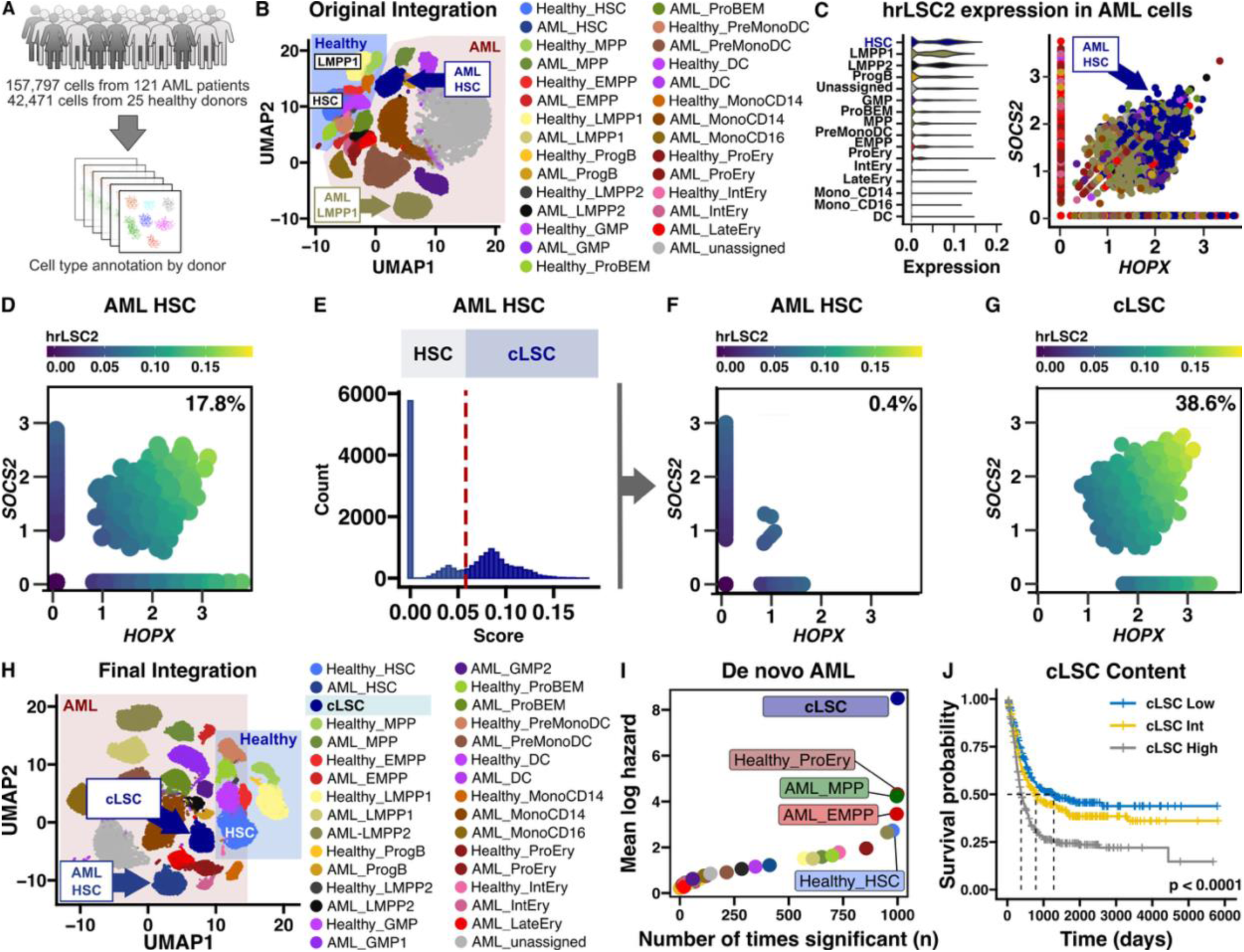
cLSCs are the most prognostic characterized cell type in de novo AML. A) Workflow for constructing an integrated scRNA-seq dataset for de novo AML. B) Post integration UMAP projection of cells from AML patients and healthy donors annotated using an HSPC-specific classifier. C) hrLSC2 expression in AML cells based on normal cell type annotations. D) Expression of *HOPX* and *SOCS2* for AML cells annotated as HSCs. Percentage reflects cells that co-express *HOPX* and *SOCS2*. E) Histogram illustrating hrLSC2 expression for AML cells annotated as HSCs. Dashed red line indicates threshold for annotating cLSCs. F-G) Expression of *HOPX* and *SOCS2* for cells re-annotated based on hrLSC2 expression. H) UMAP projection of the final integration after updating cell type annotations for hrLSC2 expression. I) Results of iterative Cox regression analysis (*n*=1000) in de novo AML evaluating the performance of cell type content for predicting survival. J) Kaplan-Meier survival analysis evaluating cLSC content. Statistical significance was tested using either the Wald or Log-rank test.

Using the updated annotations, the data were re-integrated using scGen (Figure 2H, S4F), which was then used to construct a signature matrix for bulk gene expression deconvolution using CIBERSORTx^49^. The data were split into training and test datasets, and the signature matrix was constructed using the training data. To evaluate deconvolution performance, we generated artificial bulk transcriptomes using either a single population (pure) or mixture of cell types (synthetic mixtures) from single cells in the test dataset. We observed robust deconvolution performance using our high-resolution signature matrix (*n*=32 cell types) on mixtures derived from the test dataset (Figure S4G-H) which motivated us to perform a large-scale deconvolution analysis on our cohort of de novo AML (Table ST1). The deconvolution results show balanced cell type imputations across each cohort (Figure S4I). We then evaluated the association between cell type frequencies and survival using ensemble machine learning methods (Table ST8) and observed that cLSC content was the most important feature for predicting survival. Importantly, other cell types including hrLSC2 high LMPP1s (cLSC-LMPP1) were weakly associated with outcome, highlighting that subsetting on the most immature cell subset derived from scRNA-seq, i.e. the HSC, increases prognostic power. The strength of this association is illustrated in Figure 2I which presents the results from iterative coxph regressions (*n*=1,000) for each cell type, and Figure 2J which shows how cLSC content can effectively stratify patient survival outcomes.

We further evaluated the clinical relevance of cLSCs by performing a multi-variate Cox analysis which included LSC signatures, age, sex, *FLT3*-ITD status, *NPM1* mutation status, and cytogenetic risk category (Figure 3A). We find that cLSC content is significantly associated with worse outcomes and interacts with mutation status and cytogenetics for predicting survival (Figure 3B-D). Since shorter survival in AML is often related to relapsed disease, we next evaluated changes in cLSC content at relapse. To address this goal, we compiled an AML gene expression cohort containing paired cases from diagnosis and relapse after either cytotoxic chemotherapy (*n*=47 cases) or allogeneic hematopoietic cell transplant (HCT; *n*=26 cases)^36,50–54^. Patients who receive chemotherapy alone are generally considered to have lower risk disease compared to those who receive HCT^55^. After imputing cell type frequencies using CIBERSORTx, we observed a significant increase in cLSC content at relapse post chemotherapy (Figure 3E). Although we do not see an increase in cLSC content after HCT (Figure 3F), there is significantly greater cLSC content at diagnosis in AML cases that receive HCT compared to those that receive chemotherapy (Figure 3G), and no difference in cLSC content after relapse comparing either treatment modality (Figure 3H). To further characterize the relevance of cLSCs after treatment, we compared changes in cLSC frequency to those of all other cell types (Figure 3I). For patients receiving chemotherapy, the change in cLSC content was not only the most significant but was also associated with the largest effect size (Cohen’s *d*=0.85). In contrast to cLSCs, the other prognostic cell types identified by survival analysis (Table ST8) showed minimal changes at relapse (Figure 3J). Overall, these findings demonstrate that cLSCs are both the highest hrLSC2 expressing cells and the most prognostic cell type in de novo AML.

**Figure S3:**
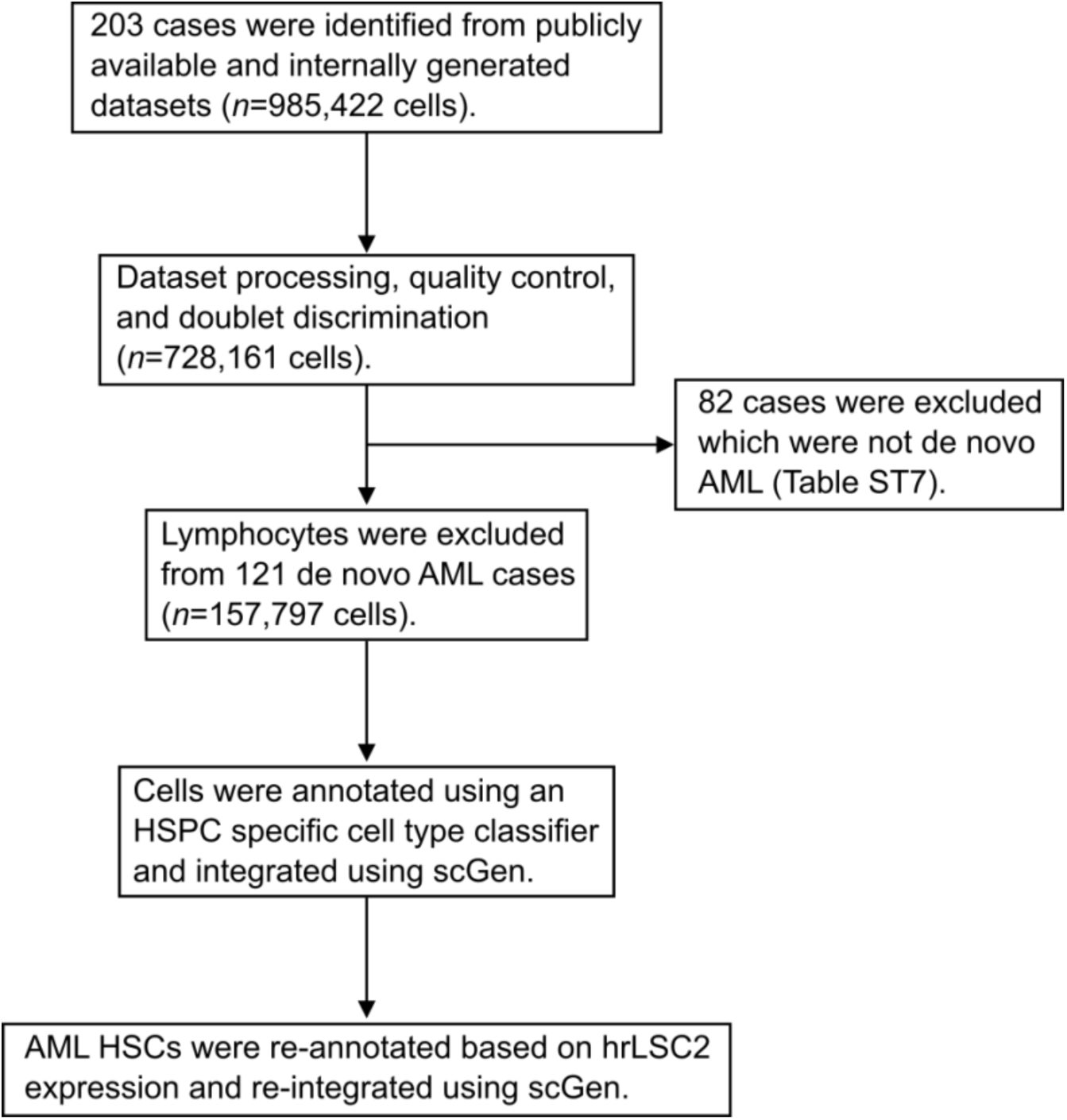
Single cell RNA-seq dataset curation, processing, annotation, and integration workflow.

**Figure S4:**
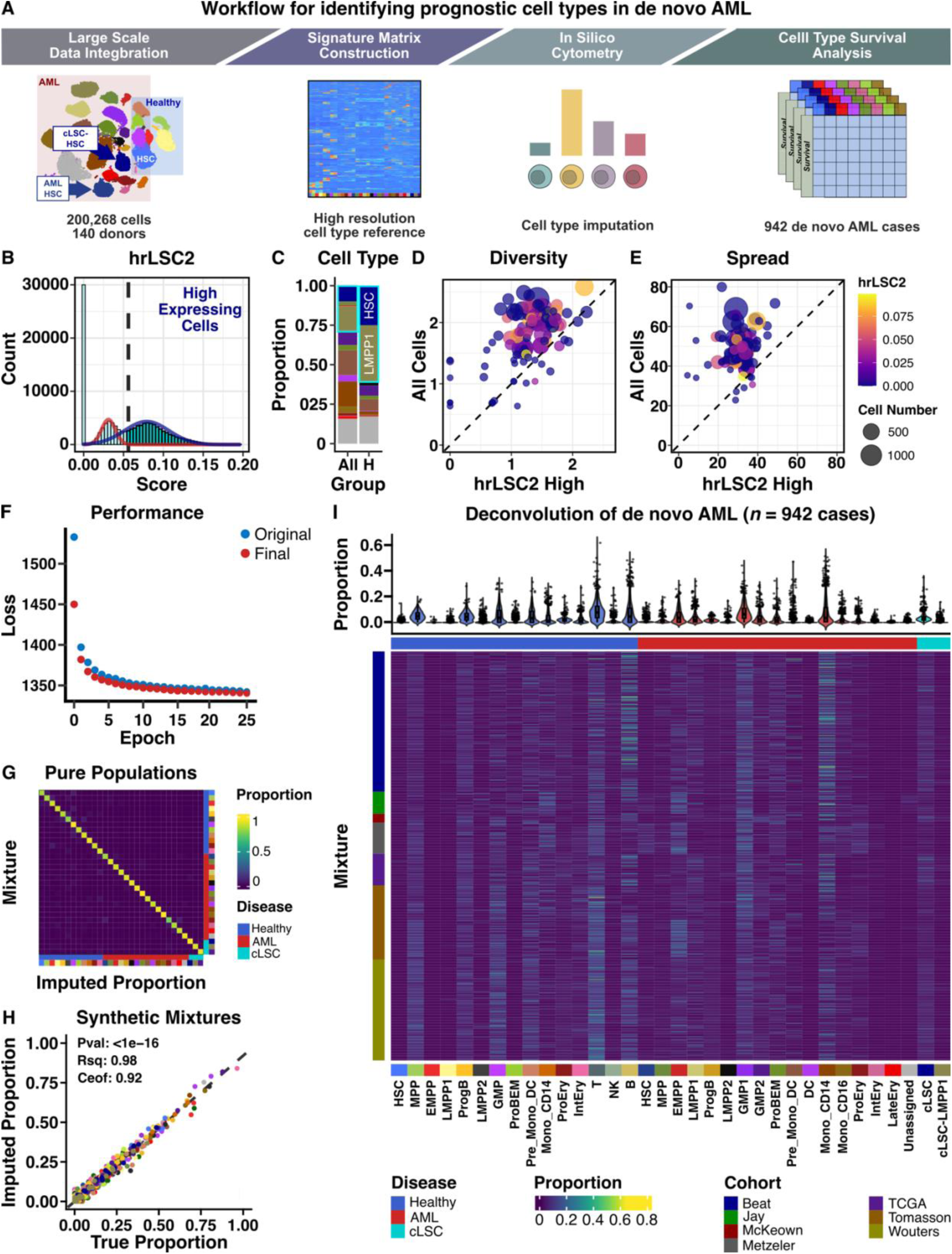
Constructing a high-resolution single cell reference for bulk gene expression deconvolution. A) Workflow for constructing a high-resolution signature matrix for a cell type specific survival analysis in de novo AML. B) Histogram of hrLSC2 expression across all AML cells. Dashed blue line indicates threshold for identifying cells expressing high levels of hrLSC2. C) Cell type frequencies within all cells and the hrLSC2 high (H) sub-population. Comparison of (D) Shannon diversity indices and (E) spread by sample for the hrLSC2 high sub-population and all cell types. The comparison containing all cells was down sampled to ensure the same number of cells. Each dot represents a single donor and is colored based on median hrLSC2 expression and sized based on cell number. F) Loss comparison over training epochs for both the original and final integration with scGen. G) Heatmap illustrating imputation results from deconvolving artificial bulk transcriptomes derived only from cells in each respective cluster. H) Comparison between imputation results and true cell type proportions from deconvolving artificial bulk transcriptomes containing mixtures of cell types at known frequences. I) Deconvolution results of 942 de novo AML cases using CIBERSORTx. The violin plot presents cell type frequencies, and the heatmap is colored based on cohort (rows) and disease (columns).

**Figure 3:**
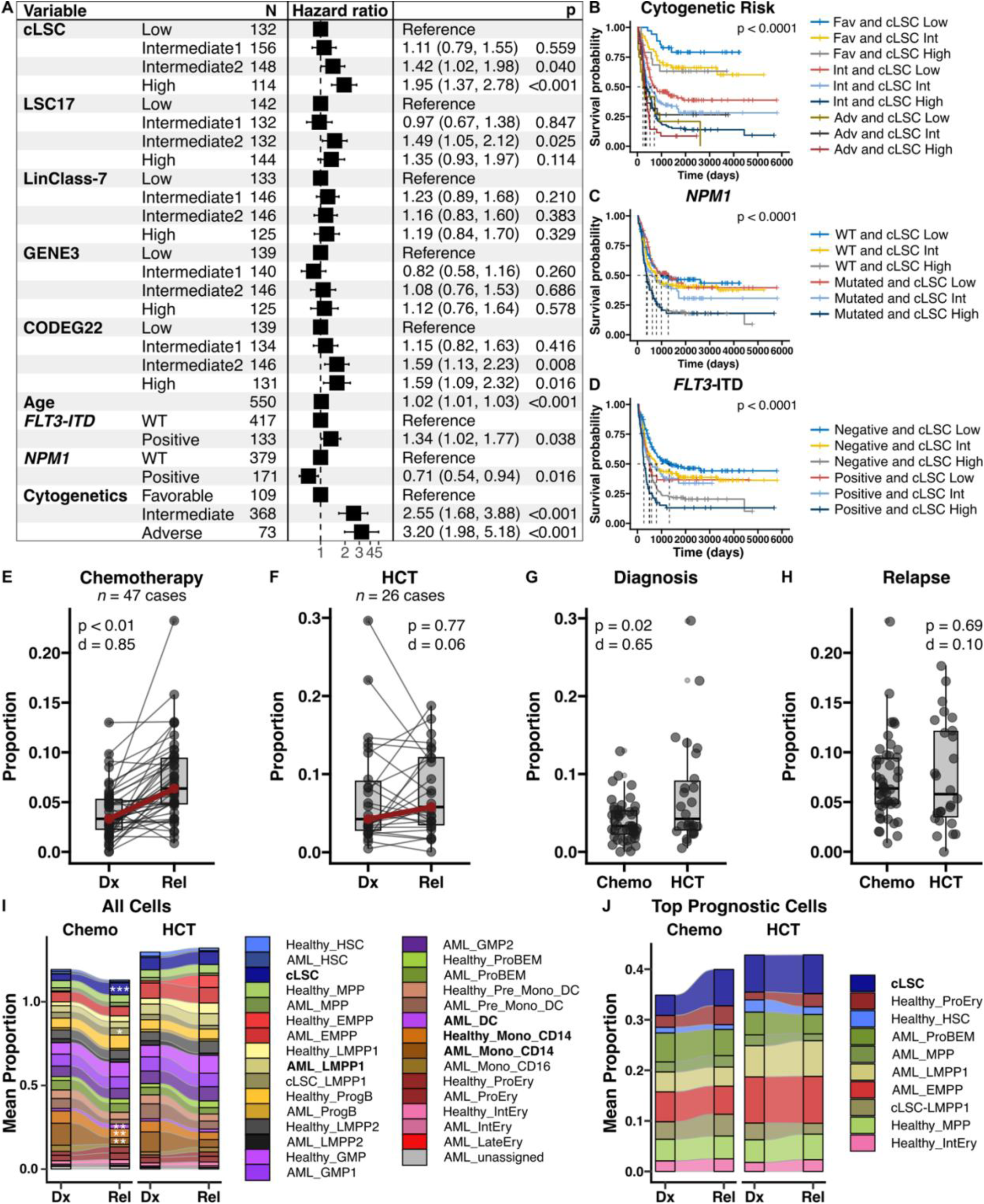
cLSCs are associated with worse outcomes in de novo AML and expand in relapsed disease. A) Multi-variate coxph survival analysis evaluating the association between cLSC content, prognostic AML signatures, age, high and low risk mutation status, and cytogenetic risk categories. B-D) Kaplan-Meier survival analysis evaluating cLSC content in combination with established high and low risk features. cLSC content at diagnosis and relapse after (E) chemotherapy and (F) HCT. cLSC content for patients receiving chemotherapy or HCT at (G) diagnosis and (H) relapse. Red line highlights median values. P-values and effect sizes (Cohen’s *d*) are indicated. I) Mean cell type proportions at diagnosis and relapse for patients receiving either chemotherapy or HCT. J) Mean proportion of top prognostic cell populations at diagnosis and relapse for patients receiving either chemotherapy or HCT. Statistical significance was tested using either the Wald or Log-rank test for panels A-D. Statistical significance was tested using t-test with multiple hypothesis correction for panels E-J. ***Adjust p-value < 0.0001, **Adjusted p-value < 0.001, *adjusted p-value < 0.05. Chemo – chemotherapy, Dx – diagnosis, HCT – hematopoietic cell transplant, Rel – relapse.

### Cells that express hrLSC2 at high levels can be identified using a unique immunophenotype

Since hrLSC2 was derived from gene sets associated with LSC content (Figure 2C), we hypothesized that cLSCs are bona fide, functional LSCs as they express the highest level of hrLSC2. To test this hypothesis, we first evaluated if these cells also express a unique immunophenotype with which they could be prospectively isolated with FACS. For this purpose, we performed scRNA-seq using a targeted mRNA panel (*n*=500) and scADT-seq (*n*=39) on 142,628 cells across 8 primary adult AML samples (Table ST5; Figure 4A). Since LSCs are often a rare sub-population, we performed a targeted mRNA analysis to efficiently profile more cells per sample compared to a whole transcriptome analysis and therefore used cells with high expression of hrLSC2 as a surrogate for cLSCs. After strict quality control, doublet discrimination, and lymphocyte, erythrocyte, and monocyte removal (Table ST6)^46^, we identified 94,989 high quality cells for subsequent analysis. hrLSC2 scores were derived per cell based on expression of *HOPX* and *SOCS2* and cells above the seventy-fifth percentile were marked as cLSCs (Figure S5). The threshold of seventy-five percent was determined based on balancing the goal of identifying cells expressing the highest level of hrLSC2 while maintaining the power to detect important features (Figure S5E). We then performed a Least Absolute Shrinkage and Selection Operator (Lasso) classification comparing ADT expression to cLSC cell status for each sample to identify protein markers compatible with FACS^56^. We observed that CD69, CD53, CD34, and CLL1 were the most important features associated with cLSC status (Figure 4B). Using the relative importance of CD69 and CD53 compared to CD34 and CLL1 as a guide, we created an iterative gating strategy for purifying cLSCs (Figure 4C-E). Using this approach, we computationally isolated a population of CD34+CLL1-CD90-CD69+CD53- cells with significantly higher hrLSC2 expression (Figure 4F) and designated this immunophenotypically defined subpopulation as iLSCs. These cells also expressed higher levels of LSC17^10^ and LinClass7^43^, two previously published LSC signatures, compared to other sub-populations^57^, albeit at a smaller fold difference than hrLSC2 (Figure 4G-H). The final CD34+CLL1-CD90-CD69+CD53- cell population was also more homogenous (Figure 4I) compared to cells purified using previously validated LSC sorting strategies such as CD34+CD38- and expression of CD99 and TIM3^57^ (Figures 4J-L).

**Figure 4:**
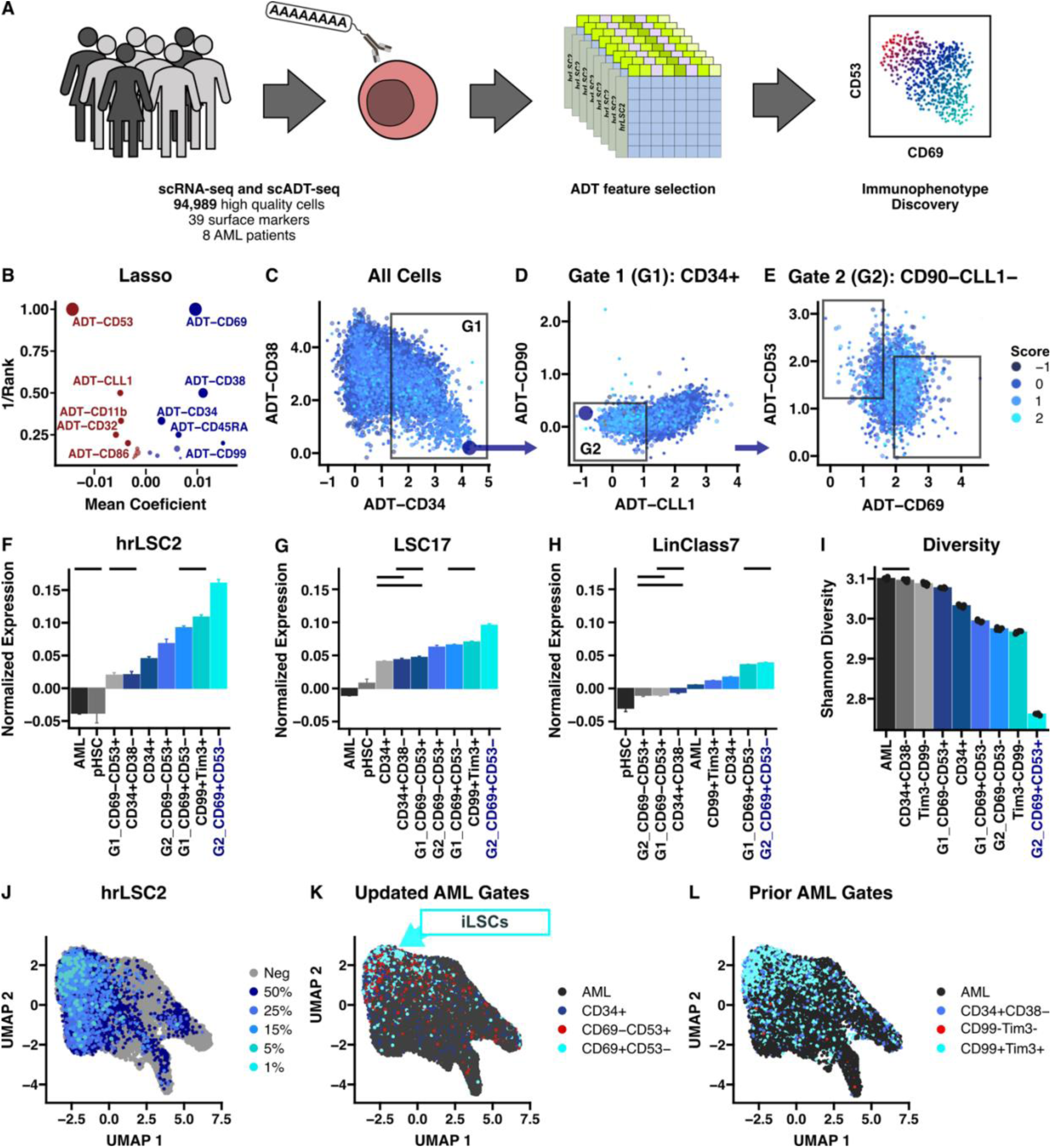
Cells expressing high hrLSC2 also express a unique immunophenotype. A) Schematic illustrating the workflow for identifying an immunophenotype for cLSCs (i.e. iLSCs) using a targeted gene expression panel and scADT-seq. B) Plot illustrating ADT rank and mean coefficient from Lasso regressions by sample comparing ADT expression and hrLSC2 status. C-E) Computational sorting strategy for purifying iLSCs. Normalized expression of F) hrLSC2, G) LSC17, and (H) LinClass7 within each computationally purified sub-population. I) Diversity measured by the Shannon diversity index for each purified sub-population. All comparisons are statistically significant unless marked by a solid line. UMAP projections illustrating (J) hrLSC2 expression or sub-population gating based on the (K) updated sort or (L) prior gating strategy. Statistical significance was tested using a Wilcoxon ranked sum test with multiple comparison adjustments with an adjusted p-value < 0.05 considered significant. scADT-seq – single cell antibody directed tag sequencing, G1 – gate 1, G2 – gate 2.

To further validate our findings, we performed a similar analysis using the updated sorting strategy on 4 cases with concurrent whole transcriptome quantification and scADT-seq data (Table ST5) and saw similar patterns in enrichment of gene signature expression (Figure S6A-C) and purity (Figure S6D-G). We then evaluated the broader relevance of this observation by computationally purifying cells from our large scRNA-seq dataset (*n*=121 AML cases; Figure 2) using expression of the genes defining the iLSC immunophenotype, as only RNA data was available for this large number of cases. We note that cLSCs reside within the Lin-*CD34*+*CLEC12A*- gate (Figure S6H) which is comparable to the iLSC immunophenotype derived from our multi-omic analysis (Figure 4). These observations suggest that an iterative strategy is most effective for isolating a relatively pure population of cells expressing hrLSC2 at high levels, which we identify as iLSCs.

**Figure S5:**
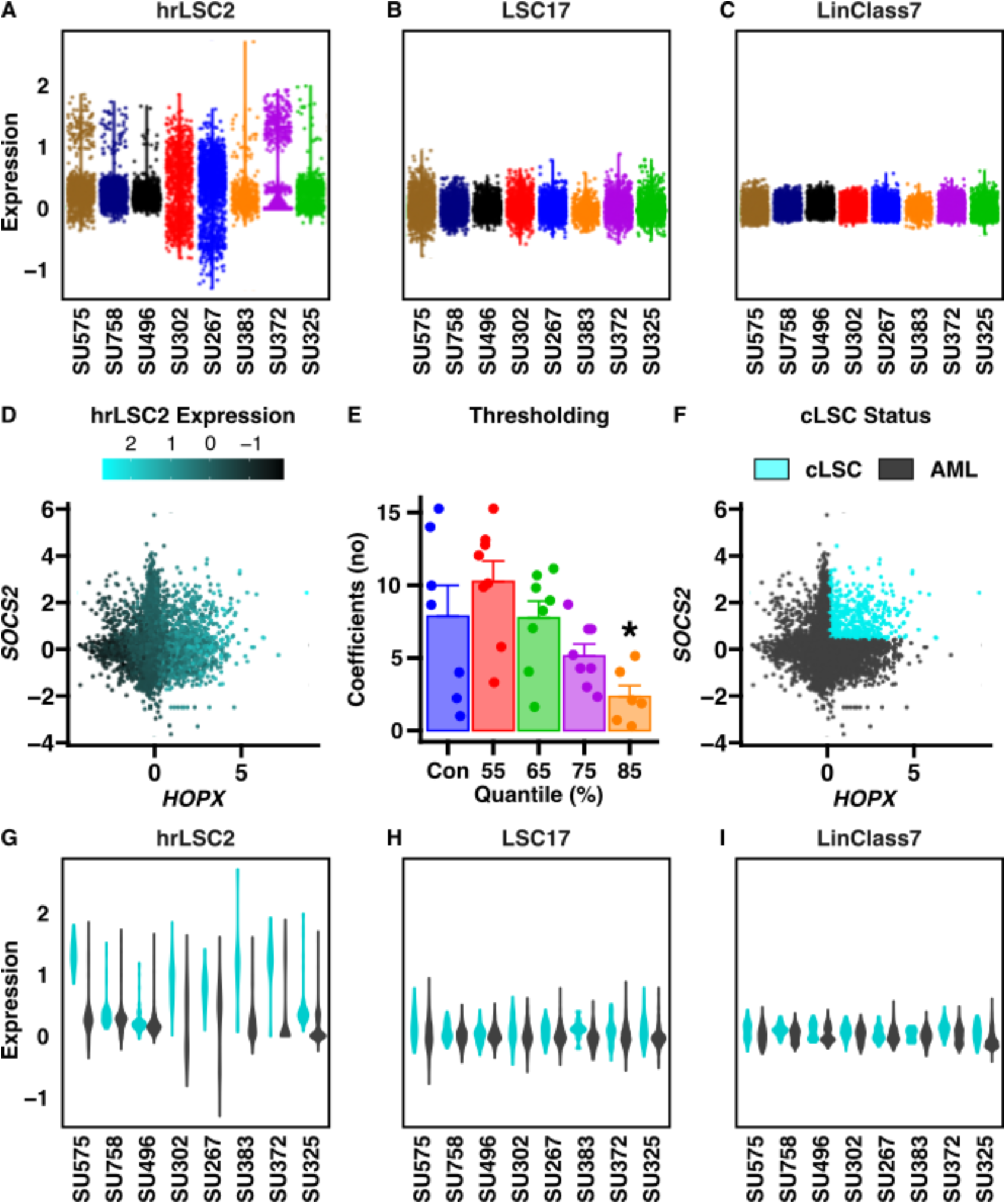
hrLSC2 expression can be used to mark a sub-population of AML cells. Cellular expression of (A) hrLSC2, (B) LSC17, and (C) LinClass7 for each donor. D) Scatter plot comparing *HOPX* and *SOCS2* expression, where each cell is colored based on its calculated hrLSC2 score. E) Number of non-zero coefficients using Lasso regression comparing ADTs and hrLSC2 as a continuous variable (Con) or using Lasso classification based on quantile thresholding for determining cLSC status. A threshold of 75% (i.e. the top 25% being considered cLSCs) was determined to be optimal as there is a significant decrease in coefficients with higher thresholding. F) Scatter plot comparing *HOPX* and *SOCS2* expression and cLSC status based on top 25% thresholding. G-I) Expression of gene signatures based on cLSC status. Statistical significance was tested using a t test or ANOVA with multiple comparison adjustments. ∗Adjusted p < 0.05.

**Figure S6:**
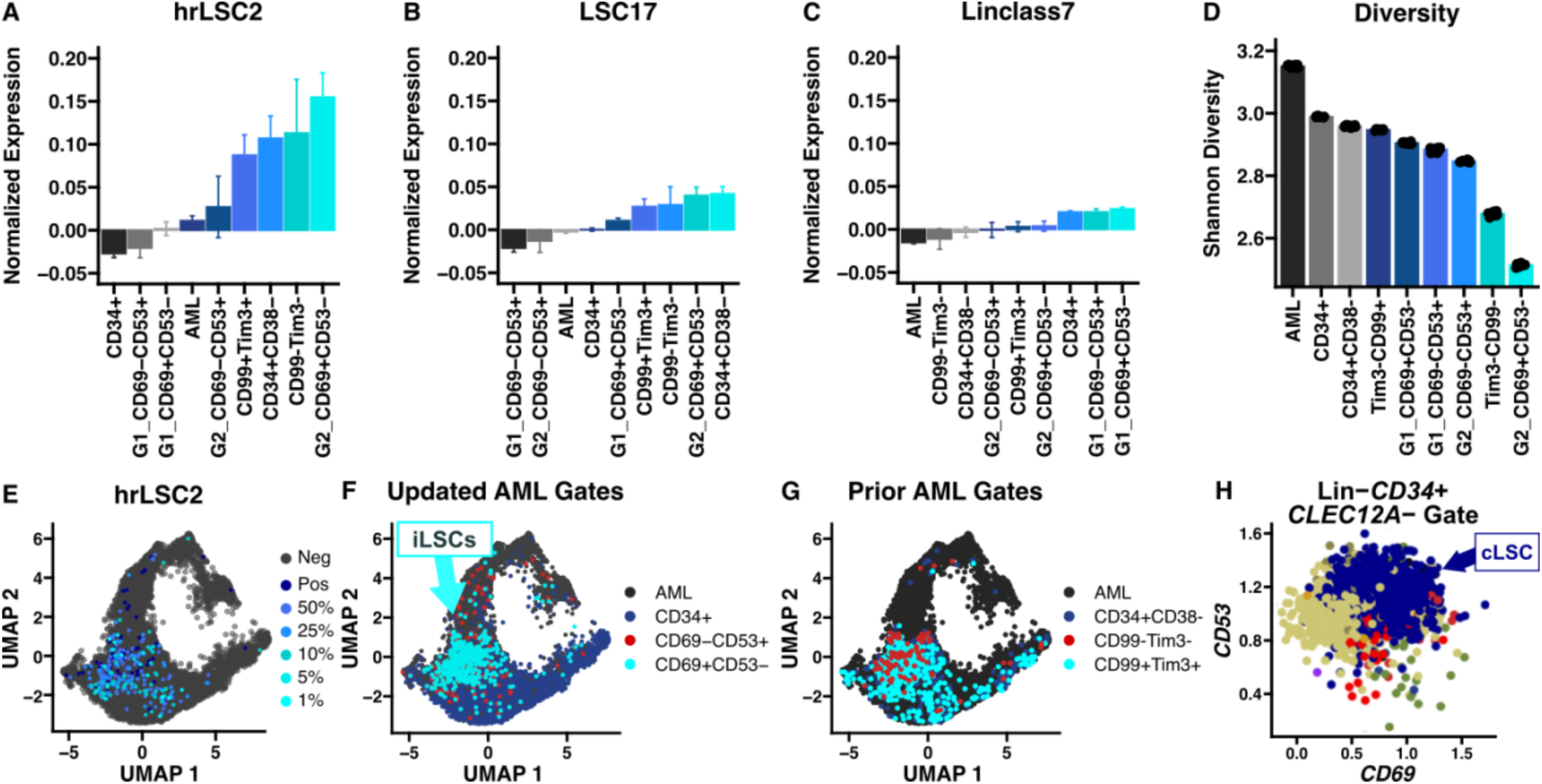
Computationally purifying iLSCs can increase sub-population hrLSC2 expression and purity. Results from purifying AML cells using whole transcriptome scRNA-seq and scADT-seq. Normalized expression of (A) hrLSC2, (B) LSC17, and (C) LinClass7 within each purified sub-population. D) Diversity measured by the Shannon diversity index for each purified sub-population. UMAP projections illustrating (E) hrLSC2 expression or sub-population gating based on the (F) updated computational defined sub-populations or (G) prior LSC gating strategies. H) *CD53* and *CD69* gene expression for cells computationally purified from the scRNA-seq cohort (Figure 2) based on negative lineage marker (Lin-), *CD34*, and *CLEC12A* (gene encoding for CLL1) expression which corresponds to G2. G1 – gate 1, G2 – gate2.

### cLSCs have distinct biomolecular programs compared to AML HSCs and Healthy HSCs

We subsequently evaluated whether cLSCs exhibit unique biological properties compared to other HSCs by performing gene set enrichment analyses (GSEA) on differentially expressed genes between cLSCs and the combination of AML-HSCs and healthy HSCs or cLSCs and healthy HSCs alone (Figure S7). We identified 307 and 265 significant gene ontology (GO) terms respectively which are presented in Figure S7A-B, and the top GO terms are presented in Figure S7C-D. The terms were clustered based on semantic similarity and labeled based on over-represented words present in the GO term descriptions. cLSCs up-regulate pathways associated with cytoskeleton organization, motility, cell activation, differentiation, and apoptosis. We also observe down-regulation of pathways associated with erythroid biology, RNA biology, and proteostasis. The analysis suggests that cLSCs differentially express several key pathways that may influence their development and pathogenesis.

### CD34+CLL1-CD90-CD69+CD53- iLSCs are enriched for LSC content

A primary goal of this study is to identify an LSC-enriched population and iLSCs appear to represent such a population as they are associated with worse outcomes in de novo AML, expand in relapsed disease, and express a unique immunophenotype. We therefore evaluated if iLSCs are enriched for LSC content, by designing a FACS-strategy based on our computational observations (Figure S8). Cells from 8 primary AML samples (Table ST5) were purified using this approach and evaluated *in vivo* for LSC content^58^ (Figure 5A). This cohort comprises primarily de novo AML samples that are diverse with respect to age, sex, and mutational status, and harbor common AML-associated mutations including those in *NPM1*, *FLT3*, and *DNMT3A*. We defined LSCs as cells that produce human CD45+HLA+CD33+CD19- engraftment in xenotransplanted mice (>0.1% human to mouse chimerism) as has been standard in the field^7^. We found that CD34+CLL1- cells engraft at high rates compared to CD34+ cells (Figure 5B). Importantly, we also observed that CD34+CLL1-CD90-CD69+CD53- cells produced significantly greater relative engraftment compared to CD34+ cells and CD34+CLL1-CD90-CD69-CD53+ cells (Figure 5C, Figure S9). We then performed an LDA using primary cells from 3 AML donors (Table ST9) and observed a significantly greater LSC frequency (13 to 162-fold increase) in CD34+CLL1-CD90-CD69+CD53- cells compared to CD34+ cells (Figure 5D-F). Overall, purifying primary AML cells using the iLSC immunophenotype significantly enriches for LSC content across a diverse panel of primary adult human AML samples.

**Figure 5:**
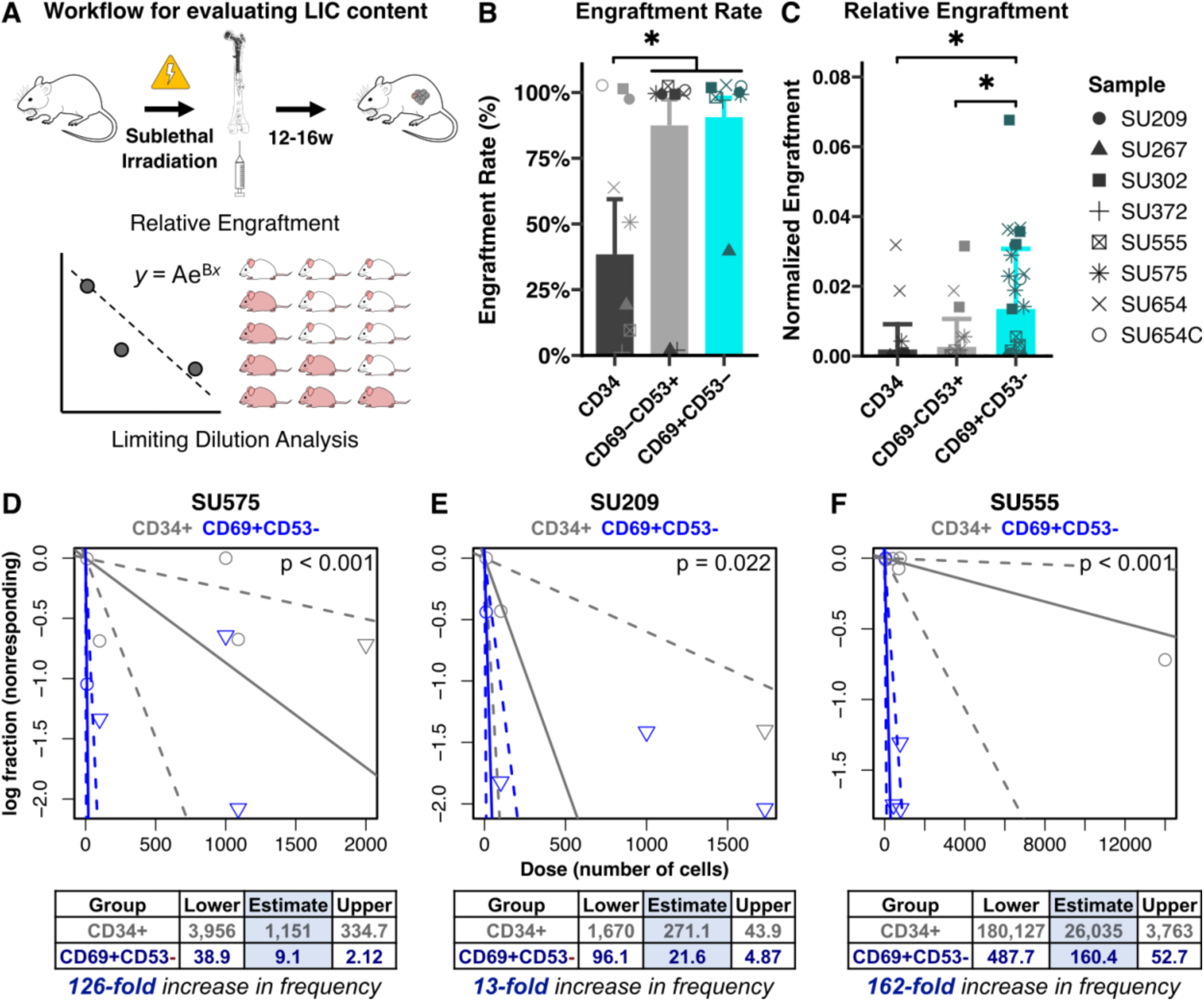
CD34+CLL1-CD90-CD69+CD53- iLSCs are enriched for LSC content. A) Schematic illustrating the workflow for xenotransplantation assays. B) Engraftment rate and C) relative engraftment between purified subsets in the primary transplants from 8 patient samples. D-F) Results from the limiting dilution analysis for 3 AML cases and significance was determined using the chi-squared test. Statistical significance for the other comparisons was tested using ANOVA with multiple comparison adjustments. ∗Adjusted p-value < 0.05.

## Discussion

Here, we report an updated LSC gene signature, hrLSC2, which is highly prognostic in de novo AML (Figure 1-3) and is significantly enriched in a sub-population of human AML cells with increased LSC content (Figure 5). Our approach for deriving hrLSC2 builds on prior work^8–10,41^ by evaluating a large and diverse cohort of 942 de novo AML cases (excluding acute promyelocytic leukemia; Table ST1) at diagnosis. By focusing the survival analysis on LSC specific genes, we identified the most clinically relevant features associated with LSC biology. The subsequent iterative evaluation using distinct machine learning methods allowed us to confidently identify a small subset of genes for model building (Table ST3-4). We used this information to create a two gene signature which was associated with worse outcomes independent of age, high and low risk mutations, cytogenetic risk categories, and validated prognostic AML signatures in a multi-variate analysis (Figure 1D). Importantly, hrLSC2 expression can further stratify patient survival within high or low risk mutation subgroups and cytogenetic risk categories (Figure 1F-H, Figure 3B-D). The strong performance of hrLSC2 highlights its potential utility as a biomarker for improving AML survival prediction and risk stratification. The added utility is likely due to orthogonal information captured by hrLSC2, which is highlighted further in our deconvolution analysis with CIBERSORTx (Figure 2-3). Using this approach, we were able to study hrLSC2 in a cell type specific context as we identified a population of AML cells, i.e. cLSCs and iLSCs, which express hrLSC2 at high levels (Figure 2), exhibit a unique immunophenotype (Figure S6H), and are highly prognostic in de novo AML (Figure 2-3). We also show that cLSCs are not only enriched in high-risk disease but also persist or expand at relapse (Figure 3E-J). Our analysis suggests that cLSCs are clinically high-risk and that aggressive treatment modalities including cytotoxic chemotherapy and HCT are ineffective at eliminating these cells. These observations are in line with the distinct yet complimentary role of the cancer stem cell model with the stochastic gene mutation model of leukemia pathogenesis^6^. Additional work will be required to prospectively validate this signature in patients receiving novel AML therapeutics.

**Figure S7:**
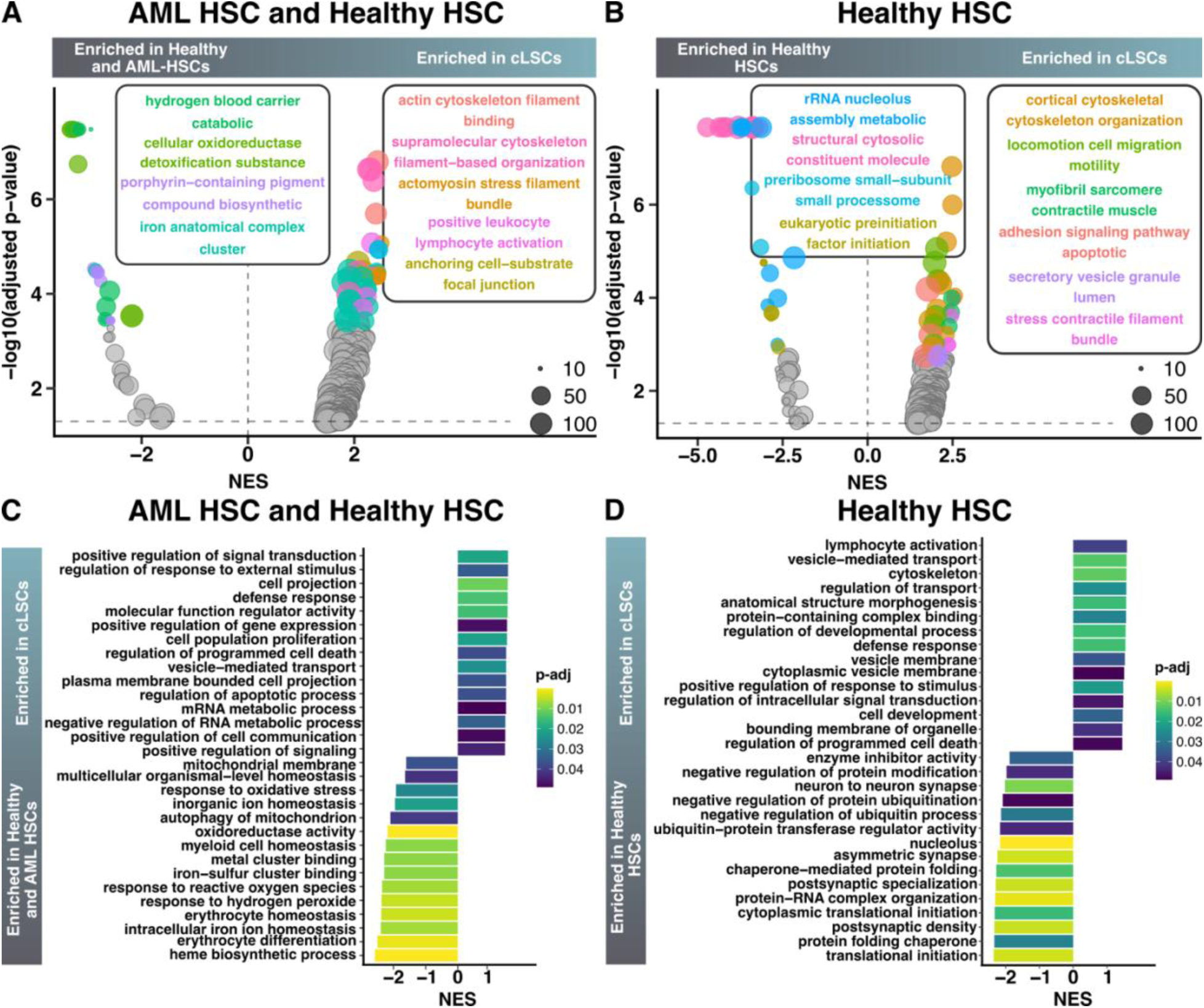
cLSCs have unique bimolecular properties. A) Volcano plot showing enriched gene ontology terms based on GSEA between cLSC and AML and Healthy HSCs. Terms were clustered based on semantic similarity, and node color indicates cluster membership. Cluster labels represent the top terms based on frequent words present in the GO term description. Node size represents gene count. B) Volcano plot showing enriched gene ontology terms based on GSEA between cLSCs and Healthy HSCs alone. The plot is formatted similarly to panel A. C) Bar plot of enriched GO terms between cLSC and AML and Healthy HSCs. D) Bar plot of enriched GO terms between cLSCs and Healthy HSCs. Colors in panels C-D indicate adjusted p-value significance levels. GO – gene ontology, GSEA – gene set enrichment analysis.

**Figure S8:**
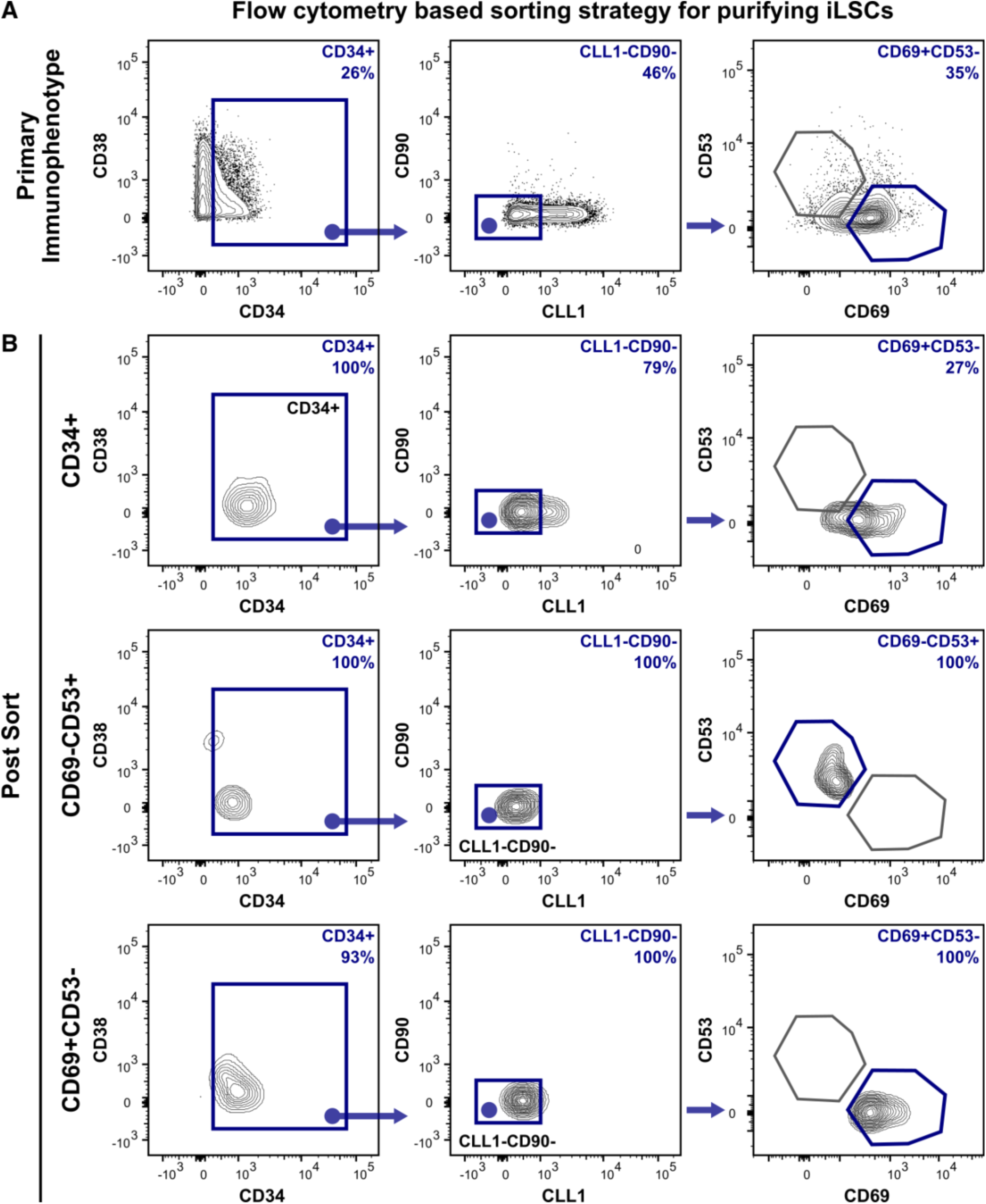
Representative plots of primary and post sort analysis of lymphocyte depleted primary AML cells. A) Sorting strategy for AML sub-populations including Lin-CD34+ AML, Lin-CD34+CLL1-CD69-CD53+, and Lin-CD34+CLL1-CD69+CD53- cells, with B) representative post sort purities for SU654.

**Figure S9:**
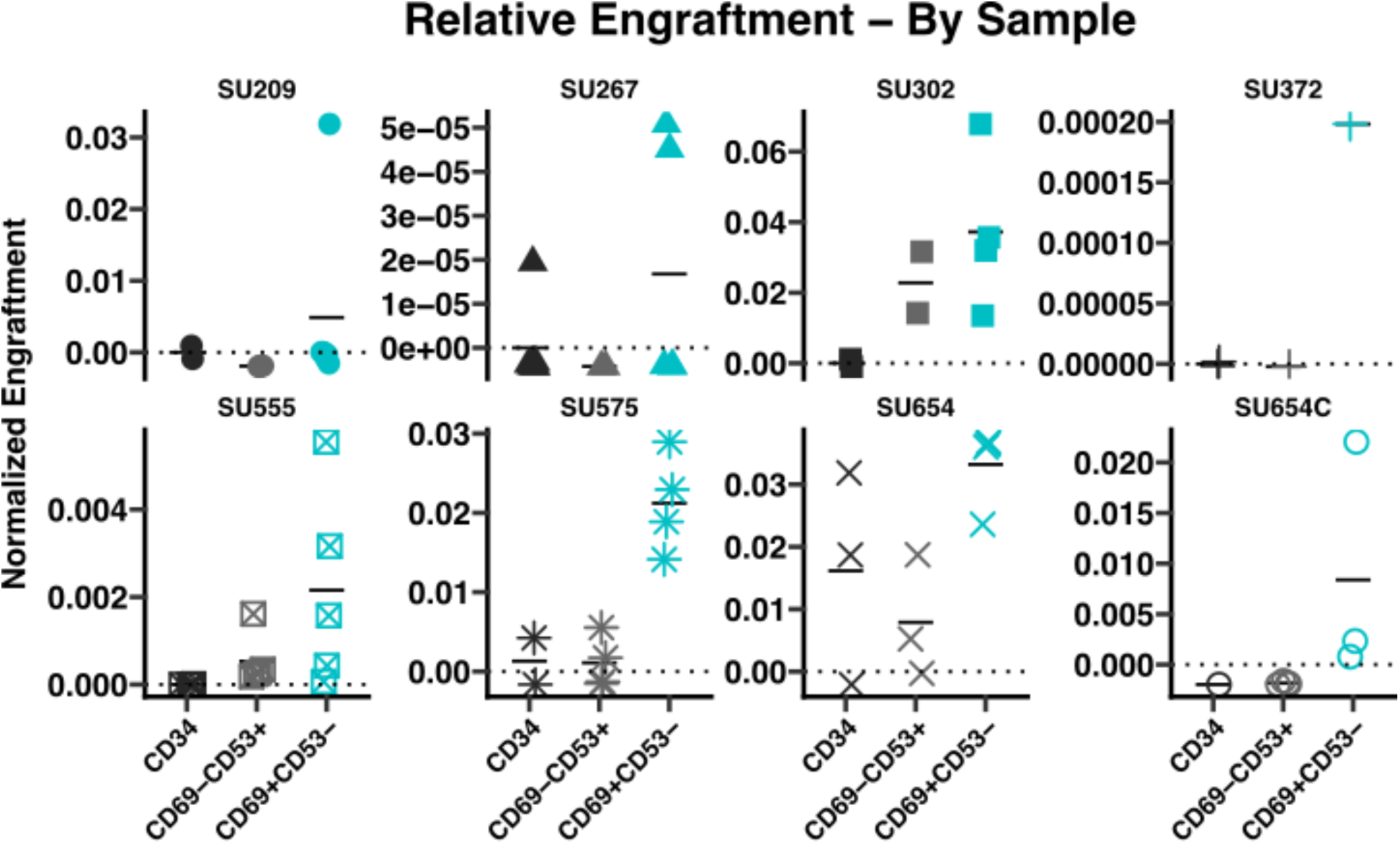
CD34+CLL1-CD90-CD69+CD53- iLSCs produce greater relative engraftment in primary transplants. The relative engraftment of purified subsets in the primary transplants for all 8 patients. Horizontal lines represent group means.

The simplicity of a two gene signature allowed us to identify cLSCs based on co-expression of *HOPX* and *SOCS2* in scRNA-seq data (Figure 4J). Importantly, cells marked by high hrLSC2 expression differentially expressed surface markers compared to the bulk AML (Figure 4B), which allowed us to identify a sub-population of CD34+CLL1-CD90-CD69+CD53- iLSCs (Figure 4F). This immunophenotype was translated into a FACS-based sorting strategy (Figure S8) and used to purify iLSCs from primary AML samples. CD34+CLL1-CD90-CD69+CD53- iLSCs consistently produced greater engraftment in the xenotransplant compared to CD34+CLL1-CD90-CD69-CD53+ cells and bulk CD34+ cells (Figure 5C, Figure S9). The results from the LDA further support that CD34+CLL1-CD90-CD69+CD53- iLSCs contain greater LSC content (Figure 5D-F). In fact, we observed an LSC frequency upwards of 1/9 in this population, which is a significant improvement over previously published flow-based LSC sorting strategies in adult human AML. The association between hrLSC2 and LSC content is important, as the ability to prospectively isolate a highly pure LSC population across a diverse collection of AML cases has previously been an unmet goal. Indeed, while previously implicated LSC markers like CD99 were associated with cLSC content, markers like CLL1, TIM3, CD123, and CD25 had variable associations with hrLSC2 (Figure 2B). Of note, we show that CLL1+ cells significantly lose long term engraftment capacity in adult AML and healthy hematopoiesis^46^, which argues against its role as a stem cell marker. The combination of hrLSC2 and the associated iLSC immunophenotype can serve as a valuable tool for identifying, tracking, and studying AML LSCs. For instance, hrLSC2 genes and surface markers can be translated into clinical assays for detecting measurable residual disease for AML patients^5,59^, which can be evaluated in future prospective clinical trials.

Although we show that *HOPX* and *SOCS2* expression can mark LSCs, the underlying biological relevance of these genes remains unclear. Hopx is a Hox family member, and expression of *HOPX* is the strongest negative predictor of clinical outcomes in AML from prior meta-analyses^8,60,61^. Prior work suggests that Hopx is both a marker of stem and progenitor cells in multiple organs^62–64^ and lymphocyte differentiation^65^. Hopx has been historically difficult to study as it does not directly bind DNA but may interact as a complex with other transcription factors (TFs) and proteins^66^. For instance, in cardiac development, Hopx has been shown to repress transcription through interactions with serum response factor and histone deacetylases (HDACs) in animal models^67^. Follow up studies have shown that Hopx interacts with HDAC2 to impede cell cycle entry and proliferation^68^. Chromatin immunoprecipitation experiments in murine hearts showed Hopx interactions in genomic regions proximal to Wnt family member genes, with Hopx inhibition leading to repressed Wnt signaling^69^. In murine hematopoiesis, Hopx disruption leads to reduced frequency of multipotent progenitors, decreased HSPC quiescence, and impaired engraftment^70^. Studies in human pluripotent stem cells revealed that *HOPX* disruption leads to a block in primitive hematopoietic differentiation^62^. Molecular profiling in these studies revealed increased activity of Wnt target genes after *HOPX* disruption. Overall, it appears that Hopx modulates stem cell quiescence and function, in part through inhibiting Wnt signaling via interactions with HDAC/TF complexes.

The second gene, *SOCS2*, is a suppressor of cytokine signaling member and has been associated with worse outcomes in AML^10,61,71,72^. *SOCS2* is upregulated upon activation of JAK/STAT and provides negative feedback in this signaling pathway. In the context of healthy hematopoiesis, the basal expression of *SOCS2* has been shown to be higher in HSCs compared to differentiated cells and is upregulated following cytokine stimulation^71^. In mice, Socs2 deficiency was not only associated with unrestricted HSPC proliferation upon stress, but it also led to decreased HSC frequency and stem cell exhaustion over serial transplantation^71^. In myeloid malignancies, JAK/STAT signaling is often dysregulated due to the underlying inflammatory milieu and activation of STAT proteins^73,74^. Additional work is needed to further characterize the role of both SOCS2 and HOPX in adult human AML LSC pathogenesis.

iLSCs also express CD69, a type II transmembrane C-type lectin receptor, which has been implicated as a stem and progenitor cell marker in human hematopoiesis. We have shown that CD69+ multipotent progenitors (MPPs) have long term engraftment capacity and can be separated from erythroid and myeloid biased MPPs in healthy adult hematopoiesis^46^. Prior work in *Mll-AF9*/*NRAS^G12V^* murine AML revealed that CD36-CD69+ cells are enriched for LSC content^75^, and follow up analysis in human AML samples showed that CD36-CD69+ cells had increased colony-forming capacity^76^. Additional studies analyzing pediatric AML samples pre and post cytotoxic chemotherapy using scRNA-seq revealed a persistent *CD69*+ HSC-like sub-population post treatment which also up-regulated *HOPX* and *SOCS2*^77^. Our analysis supports the association of CD69 with LSC content in adult human AML using xenotransplantation assays which has not been shown previously. Like the hrLSC2 genes, the mechanistic association between CD69 and LSC pathogenesis is unclear. CD69 is known to be an activation marker of lymphocytes and platelets^78^, but its role in stem cell biology is not understood. It has been shown to bind both carbohydrates and proteins, and can modulate cell metabolism, microenvironment interactions, and migration through both JAK/STAT and mTOR signaling. Although our GSEA analysis of cLSCs supports these overarching themes, future work will need to address the role of CD69 in stem cell biology.

In summary, our work provides three important advancements in the study of AML LSCs: 1) a parsimonious two-gene LSC signature, hrLSC2, which not only has a strong and independent association with poor outcomes in de novo AML but is also enriched in a specific AML sub-population, 2) an LSC computational framework for studying AML, and 3) a new immunophenotype and sorting strategy which significantly improves the purification of LSCs in adult human AML.

### Limitations of the study

This study provides significant advancements in our understanding of clinically relevant adult human AML LSCs. However, our analysis was limited by the scarcity of primary adult human AML tissue, as the frequency of LSCs in the majority of human AML is low, and the expense and effort of large-scale *in vivo* engraftment studies. Like prior studies, we had to functionally evaluate populations of cells, rather than single cells, for our xenotransplantation assays. This of course impairs our ability to definitively determine if our purified cells exist as a homogenous population or as a collection of cells with various functional capabilities even though this approach is consistent with historical advancements and refinements in LSC immunophenotyping. Regardless of these limitations, we were able to observe significant differences amongst the purified cells which will be informative when designing and implementing translational research in human AML.

## Resource Availability

### Lead contact

Further information and requests for resources should be directed to and will be fulfilled by the lead contact, Ravindra Majeti.

### Materials availability

This study did not generate new unique reagents.

### Data and Code Availability

- Single cell sequencing data have been deposited at GEO and are publicly available as of the date of publication. Accession numbers are listed in the key resources table.
- All original code used in this work is publicly available as of the date of the publication. The link is listed in the key resources table.
- Any additional information required to reanalyze the data reported in this paper is available from the lead contact upon request.

## Acknowledgements

We would like to thank Feifei Zhao and all members of the Majeti Lab for supporting this study. We also thank Bogdan Luca and members of the Gentles Lab, and Rahul Sinha and Gunsagar Gulati from the Weissman lab for their assistance and helpful discussions. This work was supported by the National Institutes of Health (NIH) under award 1R01CA251331, 1R01CA288731 and the Stanford Ludwig Center for Cancer Stem Cell Research and Medicine (R.M.). A.J.G. was supported by the National Cancer Institute (NCI) of the NIH under awards R01CA276828 and U01CA264611. A.E. was supported by the NCI under award F32CA250304, the Advanced Residency Training Program at Stanford, the American Society of Hematology (ASH) Scholar Award, and the Edward P. Evans MDS Young Investigator Award. M.H.L. was supported by the Blavatnik Family Fellowship. X.H. was supported by the ASH Medical Student Physician Scientist Award. We would like to thank Stanford University Human Immune Monitoring Center especially Xuhuai Ji, Molly Miranda, and Holden Maecker and Smita Ghanekar from BD Biosciences. Some of the computing for this project was performed on the Sherlock cluster. We would like to thank Stanford University and the Stanford Research Computing Center for providing computational resources and support that contributed to these research results. The content is solely the responsibility of the authors and does not necessarily represent the official views of the National Institutes of Health.

## Author contributions

Conceptualization, A.E., A.J.G. and R.M.; Methodology, A.E., A.J.G., A.M.N and R.M.; Investigation, A.E., Y.N., A.C.F., X.H., and B.B.; Writing – Original Draft, A.E. and R.M.; Writing – Review & Editing, A.E., A.J.G., A.M.N., and R.M.; Funding Acquisition, R.M.; Supervision, A.E., T.K., Y.K., D.K., M.H.L., A.J.G., A.M.N., and R.M.

## Key acronyms

AML: acute myeloid leukemia
cLSC: candidate leukemia stem cell
DC: dendritic cell
EMPP: erythroid-biased MPP
Ery: erythrocyte
GMP: granulocyte-monocyte progenitor
hrLSC2: two gene high risk LSC signature
HSC: hematopoietic stem cell
HSPC: hematopoietic stem and progenitor cell
iLSC: immunophenotypic LSC
LMPP: lymphoid-primed MPP
LSC: leukemia stem cell
Mono: monocyte
MPP: multipotent progenitor
ProBEM: basophil eosinophil mast cell progenitor
ProgB: B cell progenitor.

## STAR Methods

## KEY RESOURCE TABLE

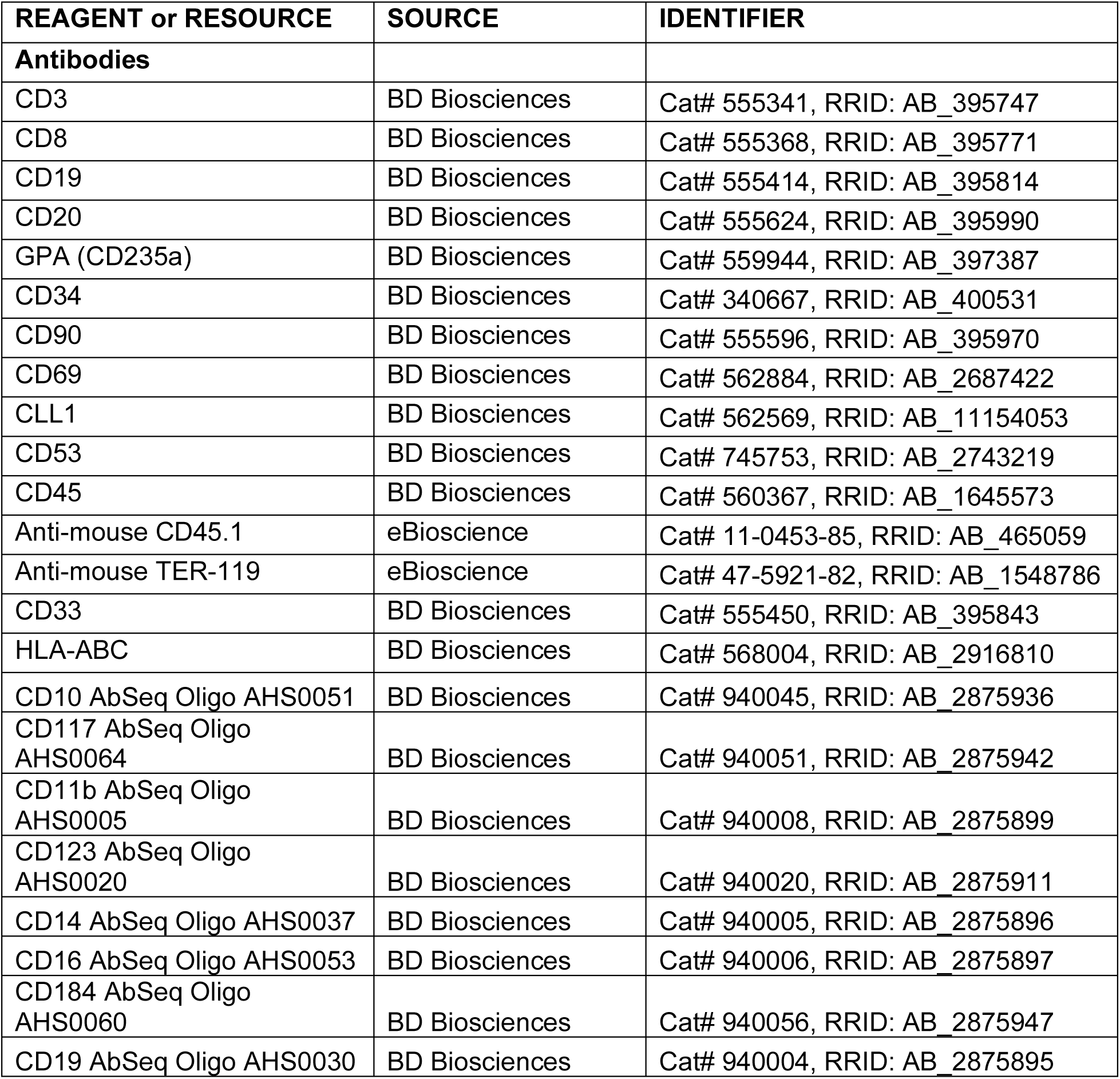
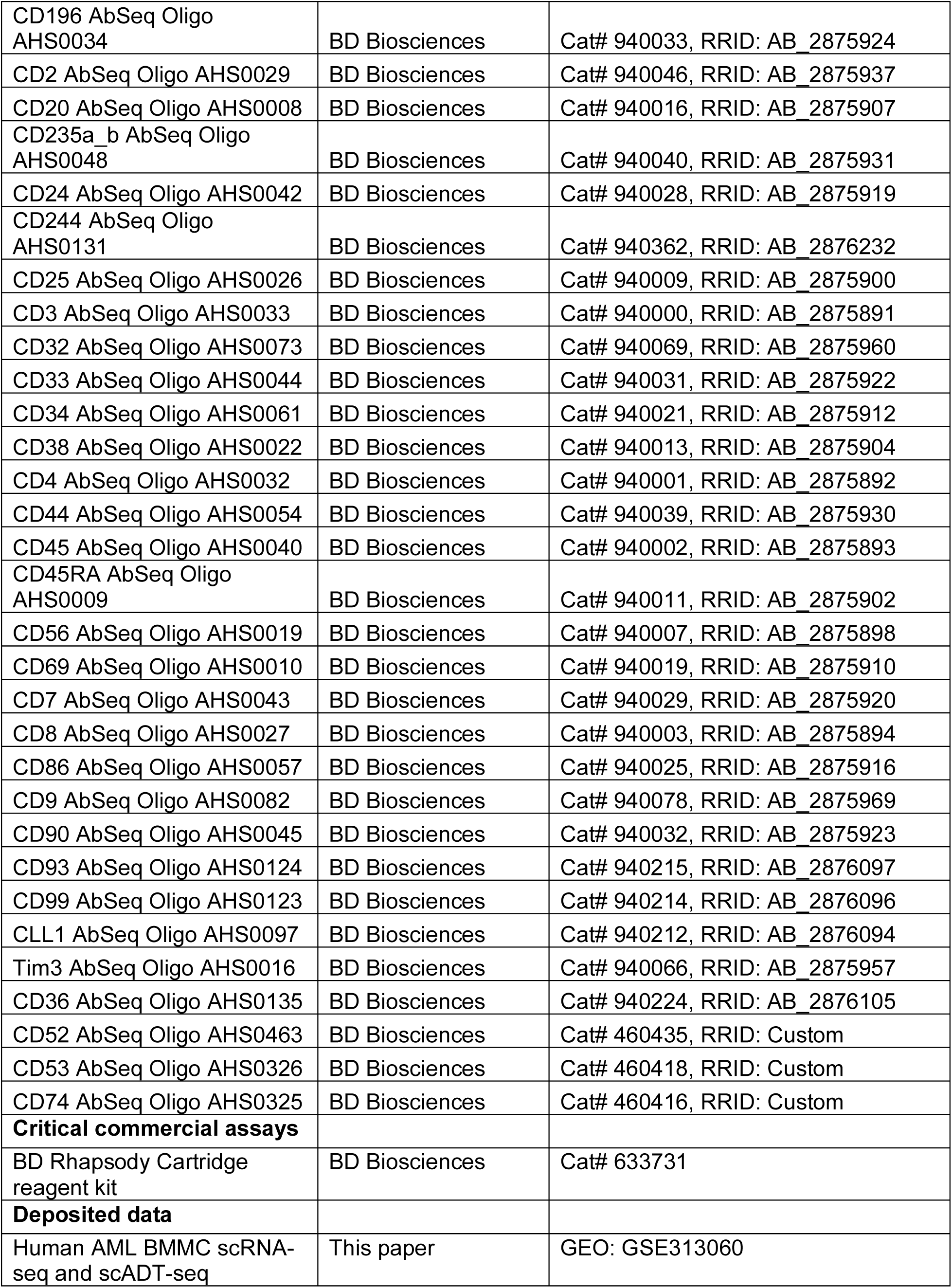
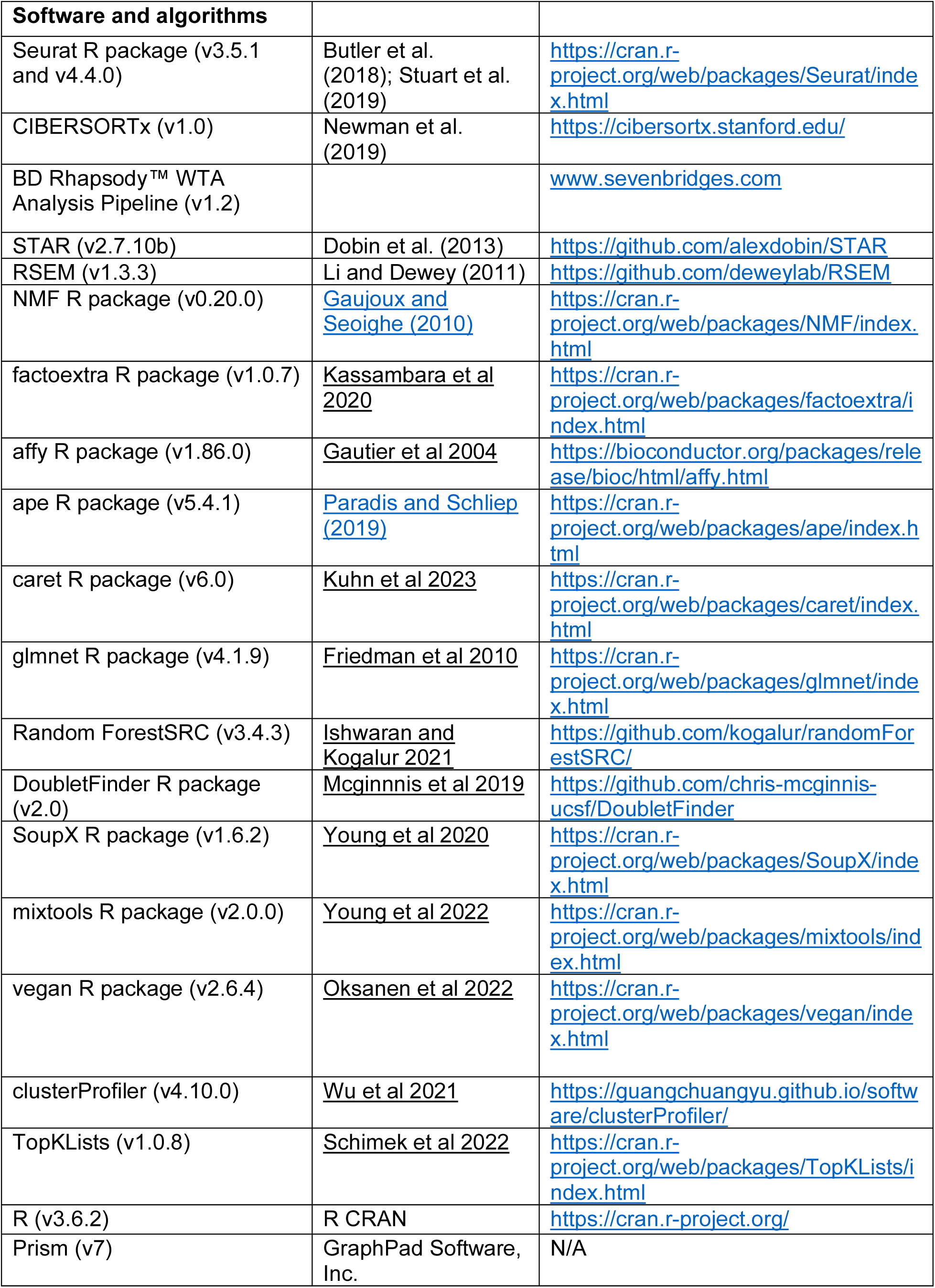
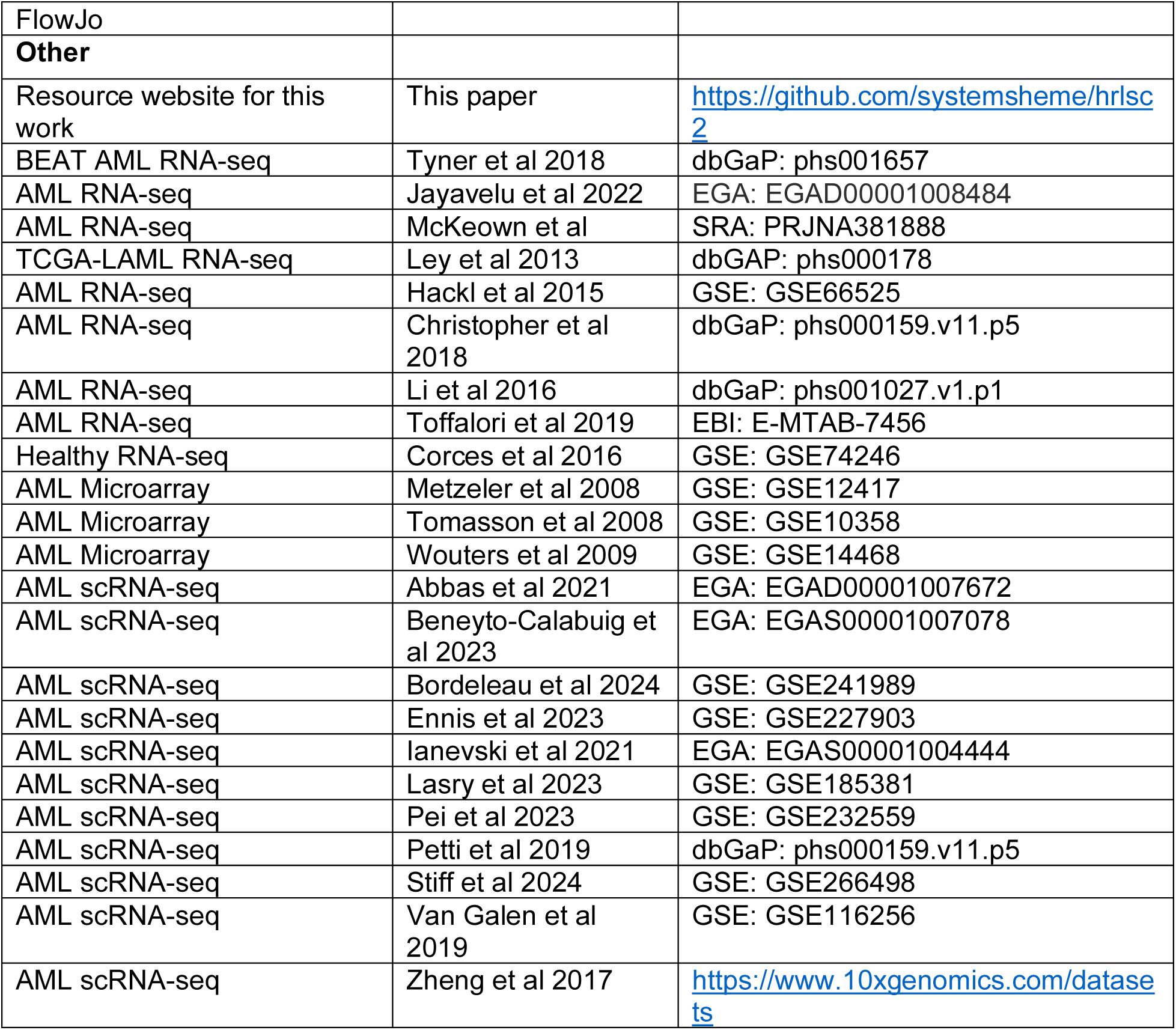

## METHOD DETAILS

### Description of healthy adult donors

Bone marrow mononuclear cells were obtained from patients with AML with informed consent and compliance with relevant ethical regulations (Stanford University IRB 6453). All cells used in this study were cryopreserved and subsequently thawed by the dropwise addition of IMDM (GIBCO, cat. no. 12440-061) containing 20% fetal bovine serum (FBS; Sigma, cat. no. F1051).

### Curation of bulk gene expression data in adult human AML

Microarray and RNA-seq data were normalized to unify data from diverse platforms using established methods^61^. For RNA-seq data, raw fastq or bam files were downloaded and aligned to the human reference genome (GRCh38) using STAR (v2.7.10b)^79^ and count tables with transcripts per million (TPM) normalization were generated using RSEM (v1.3.3)^80^. For microarray data, CEL files were downloaded and processed using the Robust Multi-array Average algorithm with the affy R package (v1.86.0)^81^. Gene counts were converted to z-values and scaled by study to facilitate cross study and platform comparisons.

### Derivation of hrLSC2

AML patients with gene expression data were filtered for those that with survival > 30 days to minimize confounding due to initial disease severity (Table ST1). We also included only de novo AML at diagnosis (excluding acute promyelocytic leukemia). Data were then split into a training and test set (2:1 split), and subset for LSC genes present in all datasets (Table ST2). Iterative univariate coxph regressions (n=1000) were performed comparing gene expression and survival. We also performed Lasso regression using the glmnet R package^82^, random forest (RF) using the randomForestSRC R package^83^, and recursive feature elimination using RF with the caret R package^84^ to identify top features. High performing features were then evaluated using ranked list analysis using TopKLists^42^. The top two features, *HOPX* and *SOCS2*, were used to create the final model and coefficients were estimated using coxph (0.4\**HOPX* + 0.2\**SOCS2*). This model was then compared to established AML gene signatures and standard clinical variables using coxph in both univariate and multi-variate models.

### Single cell captures for whole transcriptome and surface marker quantification

Cryopreserved adult bone marrow cells were thawed and stained with annexin V (Thermo Fisher Scientific) per protocol for 15 min. Cells were resuspended with DAPI and annexin V/DAPI negative cells were sorted for subsequent ADT staining and single cell capture per established protocols (BD Rhapsody™). The BD Rhapsody™ workflow is a nanowell based single cell capture system with established protocols for generating single cell whole transcription and ADT cDNA libraries. Briefly, an ADT cocktail (1uL per ADT) was prepared in a total of 100 uL of buffer. Cells were first washed and then labeled with the ADT cocktail for 30 min at 4C, washed 3 times, resuspended in sample buffer (BD Rhapsody™ Cartridge reagent kit), and captured with the BD Rhapsody™ multiomic single cell system using the manufacturer’s instructions^85^. ADT and whole transcriptome libraries were generated per manufacture’s protocol, evaluated by qubit and bioanalyzer, pooled and sequenced using NextSeq500 or Illumina Novaseq S2 (Illumina). Sequencing depth was determined based on manufacturer’s recommendations.

### Sample processing, integration, and quality control

We applied stringent quality control that incorporates the basic Seurat workflow for multimodal data processing and integration^86,87^, ambient RNA and ADT removal^88,89^, doublet discrimination using DoubletFinder^90^, and elimination of cells expressing mutually exclusive lineage markers based on prior work^46^. Fastq files generated from the BD Rhapsody™ workflow were processed on Seven Bridges (https://www.sevenbridges.com) using the standard analysis pipeline (BD Rhapsody™ WTA Analysis Pipeline) and a custom ADT fasta (Table ST10). The mRNA panel for the surface immunophenotype analysis (Figure 4, S5) was derived from differentially expressed genes from FACS sorted HSPC sub-populations, LSCs, Blasts, and normal effector cells ^57^. The top ranked genes associated with these cell types were consolidated into a 500 gene panel. The ADT panel against common surface epitopes associated with healthy hematopoietic and leukemia subsets (Table ST10). The unique molecular identifier (UMI) count matrices were imported into R (v3.6.2) and processed using Seurat (v3.5.1 and v4.4.0)^87^. Each domain, ie mRNA and ADT counts, was normalized using log-normalization or centered using log ratio normalization respectively for visualization and comparative analyses. A third concatenated dataset was created by merging normalized counts for specific down-stream analyses. Poor quality cells were removed based on total RNA content and percent mitochondrial content using the top 1% expression level. Doublet discrimination was first performed using DoubletFinder with standard metrics^90^. Doublets were removed and gene and surface marker expression were corrected for ambient noise using SoupX^88^. DoubletFinder was used to filter doublets rather than a maximum RNA content filter due to the inherent variability of RNA content and celltype^91^. Mutually exclusive lineage markers (Table ST6) were subsequently used to remove heterotypic doublets, and poor-quality cells resulting from non-specific ADT binding^46,92^. The filtered cells were subsequently integrated by sample using canonical correlation analysis (CCA) and the resulting integrated matrix was used for downstream computational analyses. For the large-scale meta-analysis, scRNA-seq count data was processed using a similar workflow (Figure S3, Table ST7). Poor quality cells were removed using sample specific mitochondrial content analysis which was generally based on the 95^th^ percentile as different single cell capture methods preferentially increase mitochondrial content. Each dataset was then labeled using an HSPC specific classifier^46^ and integrated using scGen with standard parameters and the HSPC annotation^47^.

### Single cell hrLSC2 quantification and immunophenotype derivation

Per cell hrLSC2 expression was quantified using the same coefficients described above. Cells that ranked above the top 25% expression were labeled as cLSCs and used to determine associated features based on lasso regressions. The 25% threshold was determined by performing lasso regressions using a range of thresholds. The ability to identify informative coefficients dropped significantly at thresholds higher than 25% (Figure S5). Markers were ranked based on importance derived from the lasso analysis and evaluated manually. Candidate features were subsequently evaluated using FACS to identify markers with sufficient dynamic range to efficiently purify each sub-population.

### Clustering and diversity analysis

Cells were computationally purified using the specific sorting strategies based on ADT counts. Gating thresholds were determined by modeling the expression of each ADT using mixture models (*k*=2-3) and the expectation maximation algorithm with mixtools^48^. Sub-population purity was assessed using the Shannon diversity index and the vegan R package^93^. Cellular spread was determined based on the mean Euclidean distance for each cell to the sample centroid using the principal component space.

### Deconvolution of bulk AML gene expression data using CIBERSORTx

Each AML cohort was deconvolved using an hierarchal strategy^94^ by serially applying two signature matrices. The first signature matrix was generated using bulk gene expression data from purified hematopoietic populations^57^, and was used to differentiate T, B and NK cells from HSPC, myeloid, and erythroid subsets. The second signature matrix was generated using the single cell RNA-seq data (Figure 2A) by first converting single-cell expression values to transcripts per million (TPM) space. Cells were then split into a training and test dataset using a 2:1 split by cell type. The training set was used to generate a signature matrix using CIBERSORTx using the recommended parameters^49^. Deconvolution performance was evaluated by building artificial bulk transcriptomes using the test dataset. We created pure cell type transcriptomes by combining cells based on their cell annotation, and artificial mixtures by combining cells based on randomly generated cell type frequencies based on their annotation (*n*=100). These mixtures contain cell types based on known frequencies, i.e. 1 for the pure transcriptomes or a known fraction based on the mixture. The counts from the pooled cells were converted to TPM and summed across all cells. The performance of the signature matrix was evaluated by comparing the imputed frequencies after deconvolution to the ground truth. No batch correction yielded the best results and was used for subsequent deconvolution of the bulk AML transcriptomes. The final cell type fractions were determined by multiplying the non-lymphocyte populations with the cell type content imputed by the second reference matrix.

### Gene set enrichment analysis (GSEA)

GSEA was performed by differentially expressed genes using the integrated data and the FindMarkers function and the MAST test in Seurat. Markers that were differentially expressed at an adjusted p-value of less than 0.05 were evaluated by GSEA using GO terms with clusterProfiler in R^95^. Significant terms were clustered based on semantic similarity using the treeplot function for visualization purposes (Figure 5).

### Animal care

All mouse experiments were conducted in accordance with a protocol approved by the Institutional Animal Care and Use Committee (Stanford Administrative Panel on Laboratory Animal Care #22264) and in adherence with the U.S. National Institutes of Health’s Guide for the Cand Use of Laboratory Animals.

### Fluorescence-activated cell sorting

Thawed cells were washed with FACS buffer (PBS, 2% FBS, 2 mM EDTA) and stained with antibodies for 30 minutes on ice in 50 uL total volume. Cells were then stained with Zombie Red for viability assessment for 5 minutes and subsequently washed prior to analysis. All cell sorting steps were confirmed using post-sort purity analyses. Flow cytometry was performed on a FACSAria II (Becton Dickinson). Gates were drawn by internal positive and negative controls using healthy cells and validated by back-gating on marker positive events. For CD69, gating was further evaluated using fluorescence minus one gating. Gates drawn for all other markers were consistent across all samples (Figure S7) and the panels used are listed in Table ST9.

### Xenotransplantation assays

Purified cells were transplanted into the right femur of sub-lethally irradiated NOD/SCID/IL2Rγ^-/-^-3/GM/SF (NSGS) mice (6-8 weeks, male and female, 200 rad 2-24 hours pre-transplant)^58^. Cell dosing is provided in Table ST9. Mice were harvested at 10-18 weeks by crushing the pelvis and bilateral femurs and tibias. Cells were filtered and stained using the engraftment panel (Table ST9) to assess both engraftment and lineage. Human to mouse chimerism was calculated based on hCD45+HLA-ABC+CD33+CD19- cells relative to mouse CD45+mTer119- cells. Chimerism >0.1% was considered positive for long term engraftment. For comparison across samples (Figure 5F-G), chimerism was normalized to the sum of cell dose and time (weeks). Limiting dilution analysis was performed using cell dosing presented in Table ST9 and quantified using Extreme Limiting Dilution Analysis software^96^.

### Quantification and Statistical Analysis

Details of statistical tests performed can be found in figure legends or materials and methods. For significance testing, a t-test (2-tailed distribution and two-sample equal variance) with Mann-Whitney multiple comparison adjustment was used for comparing 2 conditions or a one-way ANOVA with Tukey’s multiple comparison adjustment when comparing >2 conditions unless otherwise stated. Results with adjusted p-values <0.05 were considered significant. All statistical analyses were performed in R or Graphpad Prism.

